# Mesoscale Dynamics of Spectrin and Acto-Myosin shape Membrane Territories during Mechanoresponse

**DOI:** 10.1101/872465

**Authors:** Andrea Ghisleni, Camilla Galli, Pascale Monzo, Flora Ascione, Marc-Antoine Fardin, Giorgio Scita, Qingsen Li, Paolo Maiuri, Nils Gauthier

**Author notes:** These authors contributed equally (Andrea Ghisleni, Camilla Galli).

## Abstract

The spectrin cytoskeleton is a major component of the cell cortex. While ubiquitously expressed, its dynamic interaction with the other cortex components, including the plasma membrane or the acto-myosin cytoskeleton, is poorly understood. Here, we investigated how the spectrin cytoskeleton re-organizes spatially and dynamically under the membrane during changes in cell mechanics. We found spectrin and acto-myosin cytoskeletons to be spatially distinct but cooperating during mechanical challenges, such as cell adhesion and contraction, or compression, stretch and osmolarity fluctuations, creating a cohesive cortex supporting the plasma membrane. Actin territories control protrusions and contractile structures while spectrin territories concentrate in retractile zones and low-actin density/inter-contractile regions, acting as a fence to organize membrane trafficking events. We unveil here the existence of a dynamic interplay between acto-myosin and spectrin cytoskeletons necessary to support a mesoscale organization of the lipid bilayer into spatially-confined cortical territories during cell mechanoresponse.

## Introduction

Eukaryotic cells have developed several mechanisms to control their shape, sense their surroundings and adapt to external cues. While a lot of efforts have been devoted to the study of the acto-myosin and microtubule cytoskeletons, our understanding of cytoskeletal scaffolds directly connected to the plasma membrane (PM) lags behind. These systems are expected to play crucial roles in many cellular mechanoadaptive processes by shaping PM topology in association with the underlying cell cortex. Such processes have been investigated at nanoscale resolution through electron microscopy or advance light microscopy, and their nanoscale dynamics has just begun to be revealed by a handful of high-end microscopy techniques including fluorescence life-time, Föster resonance energy transfer, or fluorescence correlation spectroscopy (Kalappurakkal *et al*., 2019; Saka *et al*., 2014; Chugh *et al*., 2017). However, the mesoscale architectural organization and dynamics of the PM-cortex association are far less understood. In particular, we lack a detailed description of the behavior of the scaffold molecules that are part of the composite material constituted by the PM-cortex during changes in cell shape and mechanics.

A key component of this PM-cortex composite material is spectrin. This ubiquitous protein is able to assemble into a non-polarized meshwork connected to the PM, the actin cytoskeleton and their associated proteins (Bennett and Lorenzo, 2016; Machnicka *et al*., 2012). In mammals, 7 different spectrin isogenes encode for 2 α and 5 β subunits, which can be alternatively spliced into different isoforms. Among them, αII- and βII-spectrins are the most expressed in solid tissues (Machnicka *et al*., 2014; Bennett and Healy, 2009), whereas αI and βI-spectrin expression is restricted to circulating erythrocytes. Among all the spectrin isogenes, αII/βII-spectrins and αI/βI-spectrin mainly associate with the plasma membrane. At the protein level, spectrin exists as an elongated head-to-tail α/β heterodimer juxtaposed to a homologous molecule via tetramerization domains. This spectrin tetramer retains at both ends two actin-binding domains specifically harbored by the two N-termini of β-spectrin, while several PM binding domains are present along with both α and β subunits. These bonds are the key elements for anchoring the spectrin meshwork to the actin cytoskeleton and the inner leaflet of the lipid bilayer (Baines, 2009). The spectrin skeleton has been implicated in many processes, including the stability and organization of PM, signal transduction processes, and membrane trafficking via endo and exocytic pathways (Jenkins, He and Bennett, 2015). In accordance with its broad range of physiological functions, αII- and βII-spectrin genes have been found to be essential in embryonic development (Tang *et al*., 2003; Stankewich *et al*., 2011) and are also involved in many pathological conditions such as hemolytic diseases, developmental defects, cancer, channelopathies, neuropathies and cardiac defects (Lecomte, 2012).

Despite this wealth of knowledge, our understanding of spectrin macromolecular organization is limited to the study of ex-vivo erythrocytes and neurons, where it forms a triangular-like lattice and a repetitive barrel-like array interspaced by actin nodes, respectively (Liu, Derick and Palek, 1987; Byers and Branton, 1985; Pan *et al*., 2018; Xu, Zhong and Zhuang, 2013). Interestingly, erythrocytes do not possess actin filaments at their cortex. They can only polymerize short actin protofilaments made of 13 to 15 G-actin monomers (≈33±5 nm in length) that specifically serve to crosslink multiple spectrin rods, which act as the exclusive PM supportive scaffold (Ursitti and Fowler, 1994). Several attempts to describe the spectrin meshwork organization at high resolution have been reported by different electron and fluorescence light microscopy techniques. The reported lengths of the spectrin tetramer range from 50 to 200 nm, depending on erythroid or neuronal lineage and sample preparation protocols (Xu, Zhong and Zhuang, 2013; Pan *et al*., 2018; Hauser *et al*., 2018). To reconcile these disparate observations, a working model has been proposed whereby spectrin mesh can stretch and relax at maximum contour length upon mechanical perturbation to preserve PM integrity and to maintain cell shape (Machnicka *et al*., 2012). The elasticity of the meshwork is ensured at the molecular level by the intrinsic flexibility of the so-called “spectrin repeats”, triple-helix bundles that can unfold upon mechanical perturbations (Djinovic-Carugo *et al*., 2002; Law *et al*., 2003).

Its role in supporting the plasma membrane makes of spectrin a major player in cell mechanoprotection mechanisms. Recent studies in red blood cell showed that spectrin is critical in preserving cell shape by working in conjunction with myosin-dependent contractility (Smith *et al*., 2018). Whereas, in *C. elegans* neurons, spectrin protects axons from mechanical tension and deformation, in conjunction with the microtubules (Krieg *et al*., 2017). In the same model organism, spectrin and actin polymerization deficiencies have been shown to impair body axis elongation, supporting a cooperative mechanoprotective mechanism of the two cytoskeletons at the tissue scale (Lardennois *et al*., 2019). βII-spectrin has also been involved in the maintenance of epithelial cell-cell contact through microtubule-dependent processes, and its dynamics was shown to inversely correlate with endocytic capacities (Jenkins, He and Bennett, 2015). A mechanoresponsive role during myoblasts fusion in muscle development has recently been proposed for the αII/βV-spectrin dimer (Duan *et al*., 2018). This developmental process is conserved among different species (e.g. drosophila and mammalian cells), lending support to the possibility that the more ubiquitously expressed αII/βII-spectrin plays a more general and widespread role in mechanoresponsive processes.

Here, we used a wide range of mechanobiology techniques to comprehensively analyze βII-spectrin behavior during cell mechanoresponse. We found that spectrin is a major dynamic component for shaping the mesoscale-topological organization of the cell cortex upon mechanical stimuli. Specifically, spectrin complements cortical actin distribution and dynamics, while the cooperation between the two cytoskeletons ensures membrane stability during mechanical challenges, ultimately preserving cell integrity. We also unveiled a fundamental role for myosin-driven contractility in the regulation of spectrin dynamics, and how the orchestrated interplay between spectrin and PM might complement the actin-driven “pickets and fencing” mechanism in regulating membrane trafficking events, such as clathrin-mediated endocytosis.

## Results

### Spectrin and Actin define distinct but complementary plasma membrane territories in cells of different origins

Spectrin has been shown to adopt different configurations in erythrocytes and neuronal axons (Xu, Zhong and Zhuang, 2013; Fowler, 2013), while the organization in other cell types is far less accurately depicted. To fill this gap, we examined the spectrin-actin supramolecular organization in a variety of mammalian cells. We focused on βII-spectrin, the most abundant isoform among the β subunits in nucleated mammalian cells (Thul *et al*., 2017) (Figure S1 B). In mouse embryonic fibroblasts (MEFs), the two endogenous subunits (αII and βII) showed, as expected, a perfect co-localization by total internal reflection microscopy (TIRFM) (Figure S1 A). On the contrary, endogenous βII-spectrin and actin displayed a remarkable complementary pattern, which was particularly prominent along the actin stress fibers that were devoid of βII-spectrin (Figure 1 A-C). This peculiar arrangement was conserved in many other cell types, primary or immortalized, of human and murine origin, derived either from normal or pathological tissues at whole-cell (Figure S1 D), but particularly on the region adjacent to the basal PM using TIRFM (Figure S1 D and zooms in Figure S2). Specifically, βII-spectrin formed a mesh-like pattern that filled the gaps between actin cables and was completely excluded from actin-rich leading-edge structures such as lamellipodia and filopodia (Figure S2). Overall, we identified four subcellular regions of complementarity in all cell lines tested: leading-edge, stress fiber-enriched cortex, actin- or spectrin-rich membrane curvatures (Figure 1 A and Figure S2). Interestingly, actin-depleted membrane curvatures were highly enriched in spectrin and *vice versa*, suggesting that the two scaffolds might aid in shaping negatively curved PM regions.

**Figure 1.**
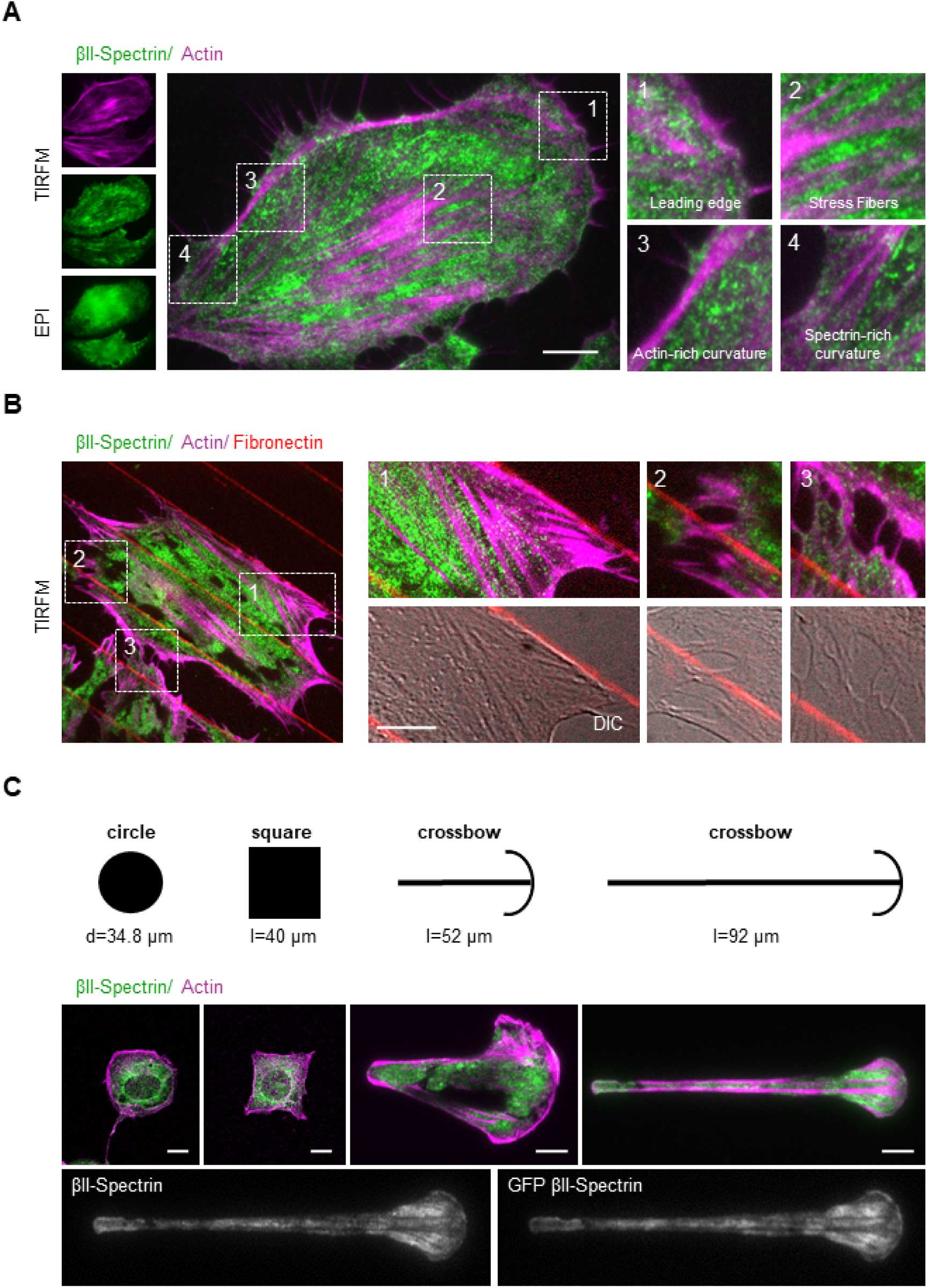
βII-Spectrin and Actin define distinct and complementary plasmamembrane territories. A) MEFs immunostained for endogenous βII-spectrin (green) and F-actin (magenta), observed by simultaneous TIRFM and EPI-fluorescence microscopy (scale bar: 10 μm). Four different cell zones are highlighted (dashed boxes, 1-4), displaying regions by distinct morphological features. B) MEFs seeded between adhesive fibronectin lines (red) and non-adhesive substrate (black), are visualized by TIRFM (endogenous βII-spectrin in green and actin in magenta, scale bar: 5 μm). Three different zones are highlighted by the dashed boxes: 1-2) cell adhesions, 3) cell-cell contact. C) Different geometries have been imposed on cells: circle and square (confocal), short and long crossbow (TIRFM). Immunostaining for endogenous βII-spectrin (green) and F-actin (magenta) are shown. The cell on the longer crossbow is transfected for GFP-βII-spectrin, immunostained for both endogenous and GFP-transfected proteins (scale bar: 10 μm).

The complementary pattern observed between spectrin and actin in cells seeded on a continuous adhesive substrate may not reflect the cortical organization of non-adhesive zones, such as on the apical part of the cell. To address this limitation, we applied microcontact printing techniques to create fibronectin-coated patterns separated by non-adhesive surface and use the resolving power of TIRFM over non-adherent membrane patches (i.e. free-standing “cortex-mimicry” zones, Figure 1 B). Also under these conditions, spectrin and actin did not colocalize, and displayed a complementary pattern at stress fiber-enriched cortex and on membrane curvatures (Figure 1 B and crossbows in C). Furthermore, by imposing different shapes to the cells from non-polarized (circle, stress fiber-poor) to polarized ones (long crossbow, stress fiber-rich), we confirmed the exclusion of spectrin from leading-edge-like zones. Finally, this distinctive distribution of spectrin and actin was also observed in fixed cells using fluorescently-tagged GFP-βII-spectrin (Figure 1 C).

### Spectrin forms a continuous dynamic meshwork of variable density during cell-driven mechanoresponses

Fibroblasts spreading can be considered as a stereotypical model to study *de novo* cytoskeletal assembly and cell-driven mechanoresponse (Figure 2 A) (Iskratsch, Wolfenson and Sheetz, 2014; Gauthier *et al*., 2009). Naïve suspended cells rapidly spread over matrix-coated substrates (fibronectin-coated glass coverslip in this work) through a multi-phasic process characterized by the initial cell attachment (P0) followed by the isotropic expansion of the cell area (P1). This expansion is propelled at the leading-edge by Arp2/3-dependent actin polymerization. After a short transition (T) driven by a change in the PM tension, the activation of myosin contractility and membrane trafficking occurs, marking the beginning of phase P2. This phase is characterized by a slower spreading rate, the maturation of focal adhesions and the formation of stress fibers (Giannone *et al*., 2007; Dubin-Thaler *et al*., 2008; Gauthier *et al*., 2011).

**Figure 2.**
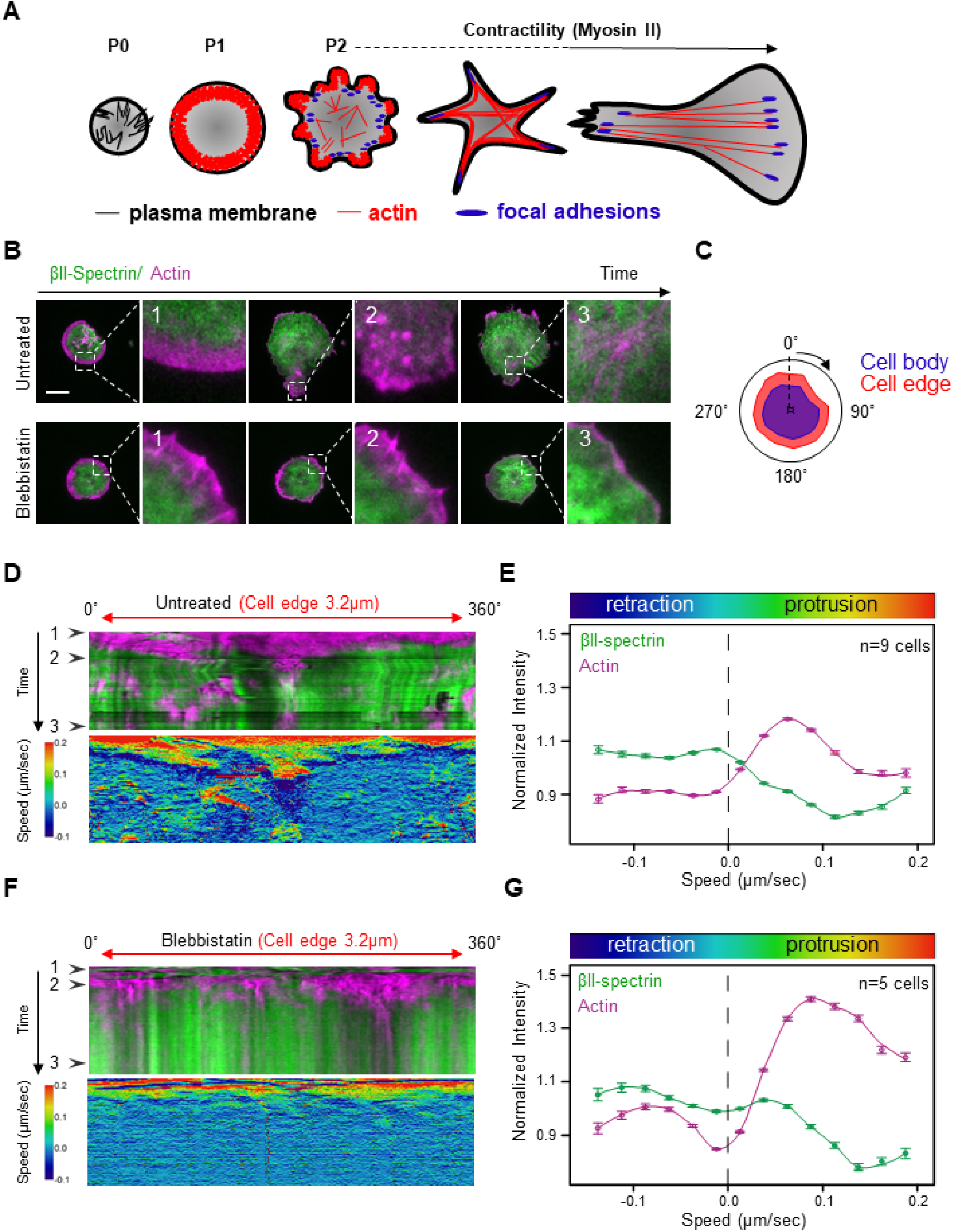
βII-Spectrin and Actin dynamics during spreading. A) Schematic representation of the different phases occurring during fibroblasts spreading on fibronectin coated surfaces. The morphological changes in terms of cell shape, actin cytoskeleton (red), focal adhesions formation (blue) and PM organization (black) are drawn. B) Cells untreated and treated with blebbistatin are visualized by live TIRFM and representative images at relevant time points are shown (green: GFP-βII-spectrin, magenta: RFP-actin, scale bar: 20 µm). Peculiar mechanisms are highlighted by white dashed boxes and are zoomed in panels 1-3. In 1 is shown a typical non-contractile phase (P1), while the contractile phase (P2) is shown in 2; in absence of myosin II-dependent contractility (blebbistatin) cell spreading is stalled. In 3, coalescent actin nodes contribute to the maturation of the actin cytoskeleton, while blebbistatin treatment impairs these dynamic processes. C) Schematic representation of the radial segmentation of the cell edge (red, 3.2 µm thickness) and cell body (blue) performed during the time-lapse. D-F) Radial kymograph analysis of cell edge behavior is presented in MEFs untreated (D) and treated with blebbistatin (F). The upper kymographs represent the integrated intensities of the two proteins (1-3 black arrowheads indicate the specific frames highlighted in the B panel), while the bottom kymographs display the edge speed related to the cell centroid. In E and G signal intensities are plotted (actin: magenta and βII-spectrin: green) in the function of speed (untreated: n=9 cells, blebbistatin: n=5 cells, mean±SEM).

To investigate spectrin recruitment to the PM during the various spreading phases, we fixed MEFs at different time points after seeding (within 5-20 minutes). We found a linear correlation between the amount of endogenous βII-spectrin and the projected cell area in the TIRF plane, likely reflecting the ability of spectrin to associate constantly with the PM (Figure S3 C-D, Table 1). Actin signal, instead, deviated more significantly from linearity as a result of a more complex and dynamic behavior during the different spreading phases, such as the transition from a lamellipodia-driven in P1 to a stress fiber-driven behavior during polarization (Iskratsch, Wolfenson and Sheetz, 2014). We also confirmed the spectrin exclusion from actin-rich protruding edges (Figure S3 A-B), in agreement with the observations at the leading-edge of polarized cells (Figure 1). However, the apparently constant spectrin/PM ratio measured at the whole-cell scale was more heterogeneous at subcellular meso-scale and evolved during spreading. In live cells, the analysis of the dynamic of GFP-βII-spectrin and RFP-actin throughout all the different phases of spreading confirmed the dynamic complementarity of the two cytoskeletons. In particular, actin was invariably associated with protrusive processes that promoted cell area growth, while spectrin displayed a passive-like behavior and was enriched in non-protrusive PM regions (Figure 2 B, Movie 1). Radial kymographs were generated to correlate fluorescence intensity in a 3.2 μm cell edge boundary with the local edge speed (see methods), where protrusions (positive speed) and retractions (negative speed) occurred over time (Figure 2 C-G). Spectrin and actin intensities displayed opposite trends. Actin intensity peaked in protruding lamellipodia (≈0.08-0.12 μm/sec) as previously observed (Dubin-Thaler *et al*., 2008), but decreased in correspondence of regions of highly positive speeds (>0.15 μm/sec) and, more significantly, when the edge movement went from null to negative speeds (Figure 2 D-E). These findings are consistent with the possibility that actin becomes diluted (less intense) in fast protruding lamellipodia (Ryan *et al*., 2012; Mueller *et al*., 2017). Spectrin intensity displayed an opposite behavior, suggesting that it may act to protect the integrity of the PM upon actin withdrawal, independent from myosin II contractility in this specific cellular compartment (see blebbistatin treatments, Figure 2 F-G). Peculiar edge-collapsing events during the contractile phase (P2), the consequence of the localized exhaustion of actin propelling activity and the subsequent actin withdrawal from the cell edge, were marked by a sudden increase in spectrin intensity (Figure S4 A-B, Movie 4). Global inhibition of contractility retained the opposite dynamic of spectrin and actin at the edge (Figure 2 F-G), but also led to increased actin polymerization during protrusion (Figure 2 G compared to 2 E). Altogether these quantitative dynamic observations provide support to a model whereby actin/spectrin are mutually exclusive both spatially and temporally along with the cell leading-edge during fast remodeling events and suggest the involvement of spectrin during cellular retraction.

**Figure 3.**
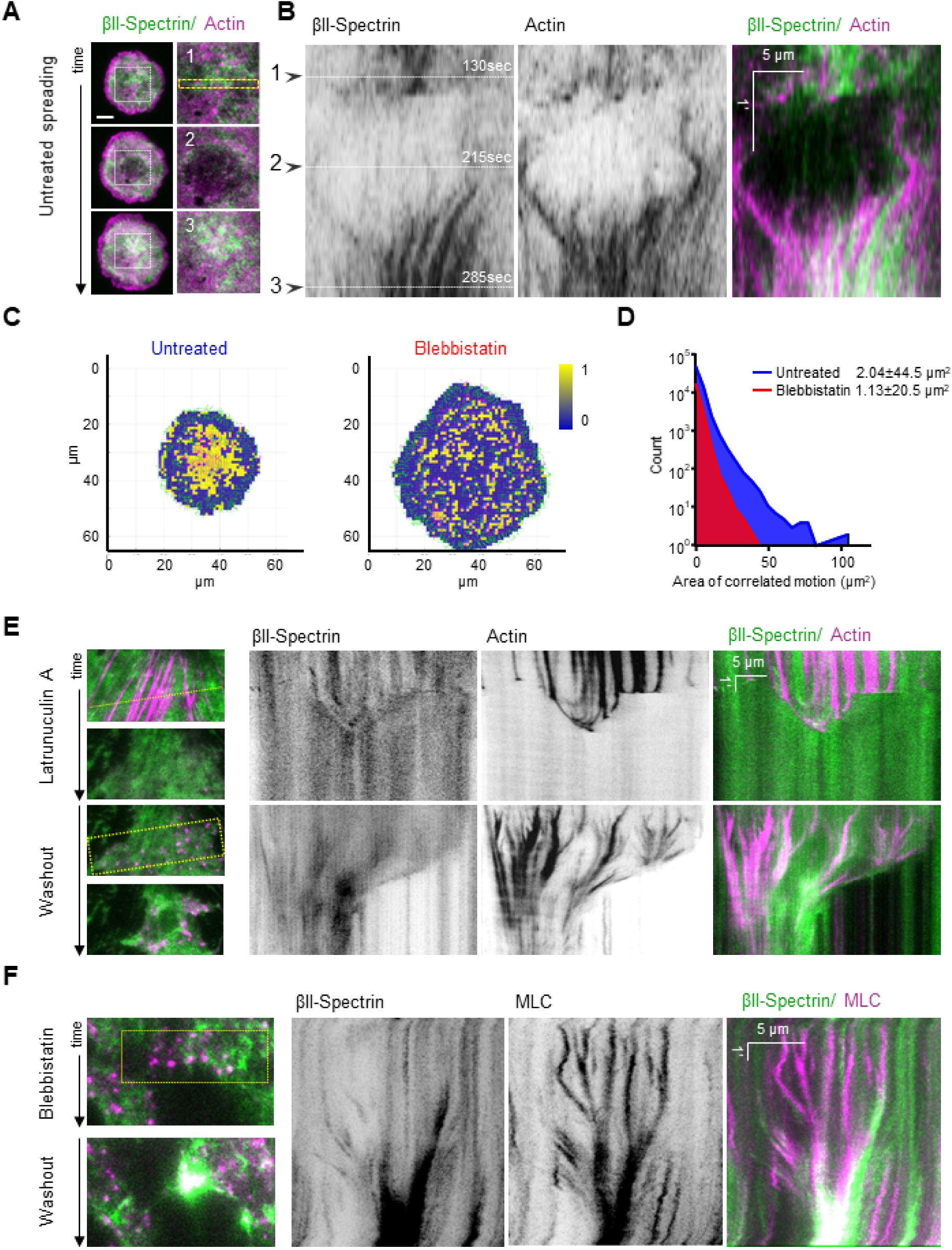
Actin nodes are instrumental for Spectrin organization. A) Cell spreading analysis at the cell body (zooms corresponding to the dashed white boxes), displayed by live TIRFM images (green: GFP-βII-spectrin, magenta: RFP-actin, scale bar: 10μm). Relevant events observed between independent experiments are shown (1-3), in particular endogenous actin nodes formation and correspondent βII-spectrin behavior. B) Kymograph generated in the region highlighted by dashed yellow rectangle. Synchronous condensation and expansion of the two proteins is highlighted by the coordinated side motion in the kymograph, despite the evident absence of colocalization, black arrowheads (1-3) indicate the respective images shown in panel. C) Two representative images of correlated PIV analyses are shown for the two experimental conditions: cells untreated and treated with 50μM blebbistatin. In yellow are shown areas of high angular and speed correlation between the two proteins, βII-spectrin and actin. Otherwise no correlation is shown in the blue zones (binary LUT). D) Total distribution of “Area of correlated motion” (yellow patches) is shown in the final graph: as predicted untreated cells (blue, n=7 cells) have higher mean area and larger distribution (±SD) compared to blebbistatin-treated cells (red, n=5 cells). E) Representative images during Latrunculin A and subsequent washout experiment visualized by live TIRFM (green: GFP-βII-spectrin, magenta: RFP-actin). Kymographs are generated in correspondence of dashed yellow line and rectangle respectively (scale and time bars are shown). F) The same experimental protocol is repeated with the drug blebbistatin and representative images are shown (green: GFP-βII-spectrin, magenta: RFP-myosin light chain). Kymograph generated in correspondence of dashed yellow box. Similar to endogenous actin nodes formation during spreading, coordinated motion is observed during the drug and washout treatments between the two channels, in absence of colocalization (time scale and scale bar are reported in the kymograph).

**Figure 4.**
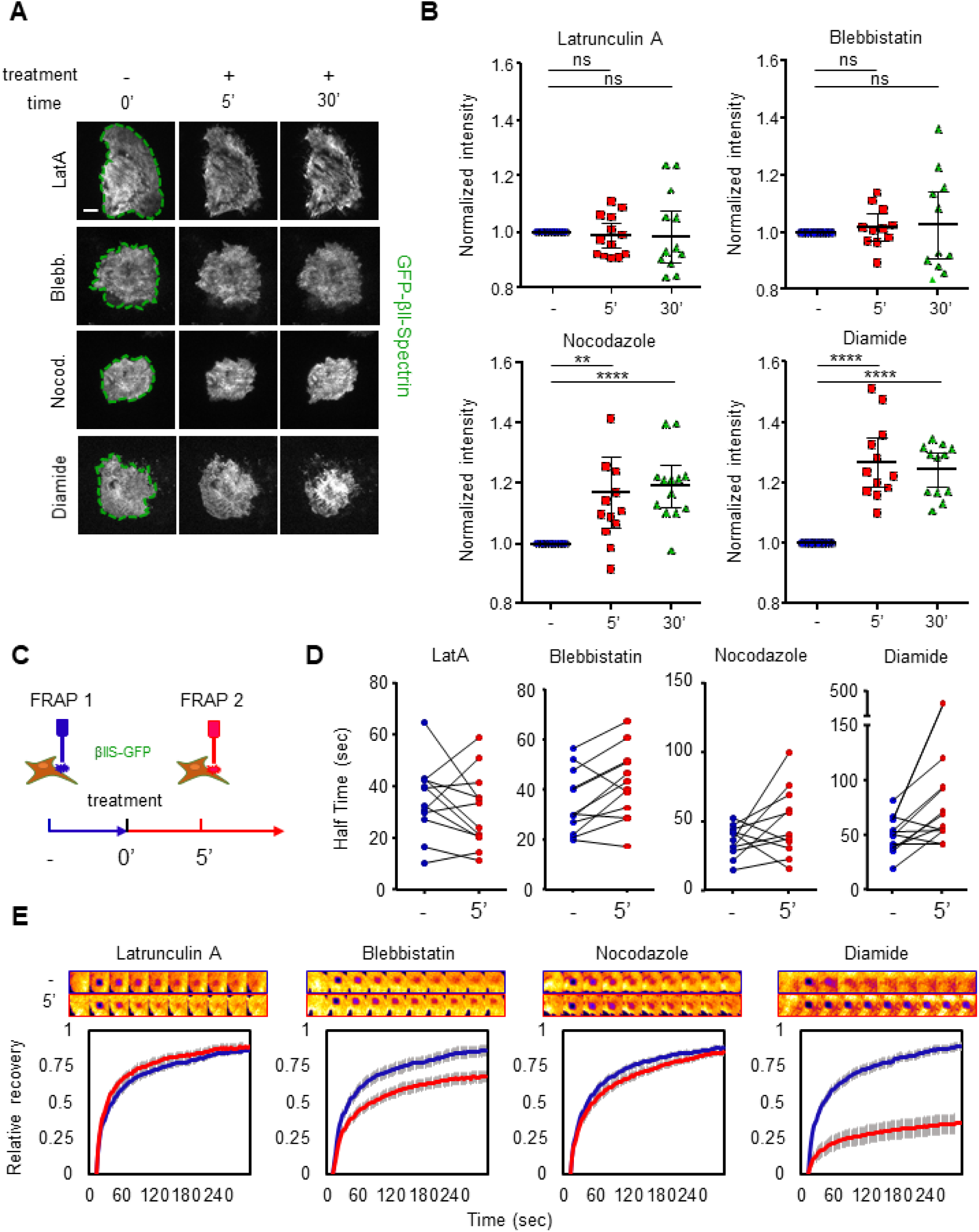
βII-Spectrin turnover relies on Myosin II-dependent contractility. A) GFP-βII-spectrin expressing MEFs imaged by live TIRFM during the administration of cytoskeletal impairing drugs are shown before (-) and during (+) the treatments (scale bar: 20 µm). Whole-cell mean fluorescence intensities are normalized to the pre-treatment frames (blue circles), and plotted in B at 5 minutes (red square) and 30 minutes (green triangle) of treatment (n=12 cells, mean±SD, paired T-test: ****p<0.0001, **p<0.01). C) Schematic representation of the dual-FRAP assay of GFP-βII-spectrin expressing MEFs, performed before (blue) and after 5 minutes of treatment (red). The resulting half-time recoveries are presented in D (individual cell connected by the black lines). Averaged half-time recoveries resulting from the single exponential fitting: latrunculin A treatment from 34.9 (-) to 27.2 (5’) seconds, blebbistatin from 34.5 to 40.1 seconds, nocodazole from 35.4 to 46.1 seconds and diamide from 38.4 to 48.2 seconds. E) Recovery curves are plotted (n=12-20 cells, mean±SEM), while the top panels show representative ROIs during the recovery phase. Mobile fractions (%) are derived from the curves: during latrunculin A treatment increased from 82.9% to 85.3%, blebbistatin decreased from 83.7% to 66.7%, nocodazole decreased from 85.1% to 82.6% and diamide treatment decreased from 85.6% to 34.5% (mean±SEM are presented in FRAP graphs to visualize the accuracy of the means subjected to the fitting procedure. See supplemental information for fitting parameters).

We next focused our attention to the spectrin dynamics under the cell body during spreading (Figure 3 A-C, Movie 2). Fixed and live TIRFM analysis showed that the spectrin mesh is progressively deployed and laid down by the cell from the back of the leading-edge during P1 (Figure 2 B and Movie 1) while apparent slight condensation in the lamella region was observable (Movie 1). Consistently, thin confocal section analysis of the dorsal cortex in P1 revealed a homogenous intermingled acto-spectrin meshwork behind the lamellipodia (Figure S3 A-B). This indicated that the non-contractile dorsal cortex of the cell was having similar organization than the ventral one. During P2, the spectrin meshwork under the cell body underwent a drastic remodeling in correspondence with the increased acto-myosin dynamic (Movies 1-2). Actin nodes were formed in this specific spreading phase, priming stress fiber maturation by condensation (Movie 2, Figure 3 A and B). Remarkably, the spectrin mesh appeared to move in coordination with these expanding and condensing nodes, albeit not showing colocalization at TIRFM resolution. Myosin II inhibition prevented such remodeling events without affecting the mutually exclusive actin/spectrin distribution at the cell edges, nor the formation of poorly mobile actin nodes in spectrin depleted zones (Figure 2 F-G, Movie 1 and 2). Cross-correlation PIV analysis of actin and spectrin flows highlighted areas of coordinated motion in terms of magnitude and directionality. This correlation landscape was analyzed during P2 in cells untreated and treated with blebbistatin, highlighting a significant decrease in size for areas of correlated motion (Figure 3 C, yellow zones) upon contractility inhibition (see the wider distribution of the measured areas in the plot of untreated cells, Figure 3 D). Thus, spectrin and acto-myosin define large membrane meso-scale territories (up to 100 μm^2^) moving in a coordinated manner, clearly highlighting that the supramolecular mesh-like organization of spectrin is dynamically cross-organized by acto-myosin remodeling.

The critical role of the acto-myosin cytoskeleton in spectrin dynamic organization was confirmed by monitoring protein flows after latrunculin A and blebbistatin washout experiments (Figure 3 E-F, Movie 3). Consistent with the physiological observation in spreading cells, spectrin expanded and redistributed upon acto-myosin stress fiber dissociation, and further augmented at cell leading-edges upon cell retraction. During the drug washout phase, acto-myosin nodes drove local spectrin coalescence as cells restored their cytoskeletal architecture (Figure 3 E-F, Figure S3 F and Movie 3). The formation of actin nodes in spectrin-less zones was also confirmed by monitoring the distribution of endogenous proteins after latrunculin A washout in free-standing “cortex mimicry” zones between patterned fibronectin lines (Figure S3 F). These results further indicate that a similar coordinated organization of the spectrin and actin meshworks occurs in the non-adhesive cell cortex.

We conclude that the spectrin cytoskeleton is a continuous meshwork tightly associated with the PM, covering it almost entirely. However, its local density under the PM can largely fluctuate upon changes in cell geometry, dynamics and mechanics. Spectrin locally condenses during events characterized by low actin-PM interaction, such as during membrane retraction at cell edges or the remodeling of cortical acto-myosin nodes that lead to the formations of actin fibers, stress fiber ultimately defining spectrin-rich territories.

### Spectrin molecular turnover depends on contractility

To address whether actin dynamics or acto-myosin contractility control the dynamics of the membrane-associated spectrin meshwork, we monitored changes in the GFP-βII-spectrin signal upon latrunculin A or blebbistatin treatments. 5 or 30 minutes after treatment, no alteration in global spectrin density (over the projected cell area) was detected by TIRFM, indicating that spectrin recruitment to the PM was independent of actin polymerization or myosin II contractility (Figure 4 A-B), in accordance with the analysis of endogenous proteins during spreading (Figure S3 C-D). We validated our approach using the spectrin/PM oxidative crosslinker diamide (Deuticke *et al*., 1983). In this case, 5 minutes after treatment spectrin intensity increased, significantly and constantly, up to 30 minutes. This result confirmed that changes in spectrin density based on fluorescence intensity could be observed upon drug perturbations by our approach (Figure 4 A-B). Since microtubules (MT) have also been proposed to control βII-spectrin and βV-spectrin dynamics at cell-cell junctions (Jenkins, He and Bennett, 2015; Duan *et al*., 2018), we investigated their role in βII-spectrin recruitment to the cell cortex. Upon MT depolymerization, even at early time points (5 minutes), spectrin density increased by almost 20% (Figure 4 A-B). Thus, MT can control, at least in part, the recruitment of spectrin to the PM. However, given the broad effects of MT depolymerization on membrane trafficking, we cannot exclude that nocodazole treatment, known to block exocytosis and not endocytosis (Gauthier *et al*., 2009), may reduce PM area driving the apparent spectrin condensation.

We next investigated the mechanisms regulating spectrin turnover by FRAP analysis upon drug treatment (LatA, Blebbistatin, Nocodazole, Diamide). Since maximal response to the drugs was observed after 5 minutes, a dual-FRAP assay on single cells expressing GFP-βII-spectrin was performed before and after 5 minutes of treatment to avoid secondary effects driven by long-term cytoskeletal perturbation (Mikulich, Kavaliauskiene and Juzenas, 2012; Signoretto *et al*., 2016) (Figure 4 C-E). The impairment of actin filament turnover by latrunculin A did not affect significantly either the half-time recovery (Figure 4 D) or the mobile fraction of spectrin (Figure 4 E, Extended Table 2). Instead, myosin-II inhibition by blebbistatin caused a significant reduction in the mobility of spectrin with an increased half-time recovery. As expected for protein crosslinking experiments, diamide-treated cells showed severely reduced spectrin dynamics. During nocodazole treatment, instead, spectrin mobile fraction was not affected, consistent with the potential indirect implication of MT in actin reorganization rather than a direct effect on spectrin dynamics. Overall, our results show that spectrin molecular turnover relies strongly on contractility.

### The actin-binding ability of spectrin is needed to coordinate spectrin dynamics with changes in cell mechanics

Since the spectrin meshwork dynamics depended on contractility and actin polymerization, we discerned the contribution of the spectrin domains that bind to actin or the membrane by deletion mutants. The actin binding domain (ABD) is present only in β-spectrins subunit which also harbors at least 3 PM anchoring points (Machnicka *et al*., 2014). We generated mutants of βII-spectrins deleted for the actin-binding (ΔABD), or the phosphatidylethanolamine-binding (ΔPE) or the phosphatidylserine-binding domain (ΔPS) (Figure 5 A-B). GFP-tagged mutants were expressed in fibroblasts, analyzed by FRAP and compared to full-length FL-βII-spectrin construct for turn-over and mobility (Figure 5 C-E, Extended Table 2 (ΔPS results are only reported in Extended Table 3)). The actin binding mutant, ΔABD, displayed an increased mobile fraction (87.2%) and a decreased half-time recovery (t_1/2_=24.56 sec) as compared to the FL construct (mobile fraction=74.8%, t_1/2_=41.7 sec). The ΔPE mutant displayed mildly decreased half-time (t_1/2_=56.8 sec) while the mobile fraction was comparable to the FL-βII-spectrin (73.2%). Similar results to ΔPE were obtained for ΔPS mutant (Extended Table 3). The individual GFP-tagged PE-domain displayed a diffusive behavior through the lipid bilayer as expected from a freely diffusive lipid-binding domain (Figure 5 D, inset). On the other hand, ΔABD, ΔPE and ΔPS mutants were all correctly targeted to the PM and excluded from actin stress fibers (Figure S4), as they likely incorporate into tetrameric complexes with endogenous αII-spectrin. These results indicate that the actin-binding domain is critical for spectrin meshwork stabilization but not its localization, while potential cooperative mechanisms exerted by different lipid-binding domains ensure PM-targeting.

**Figure 5.**
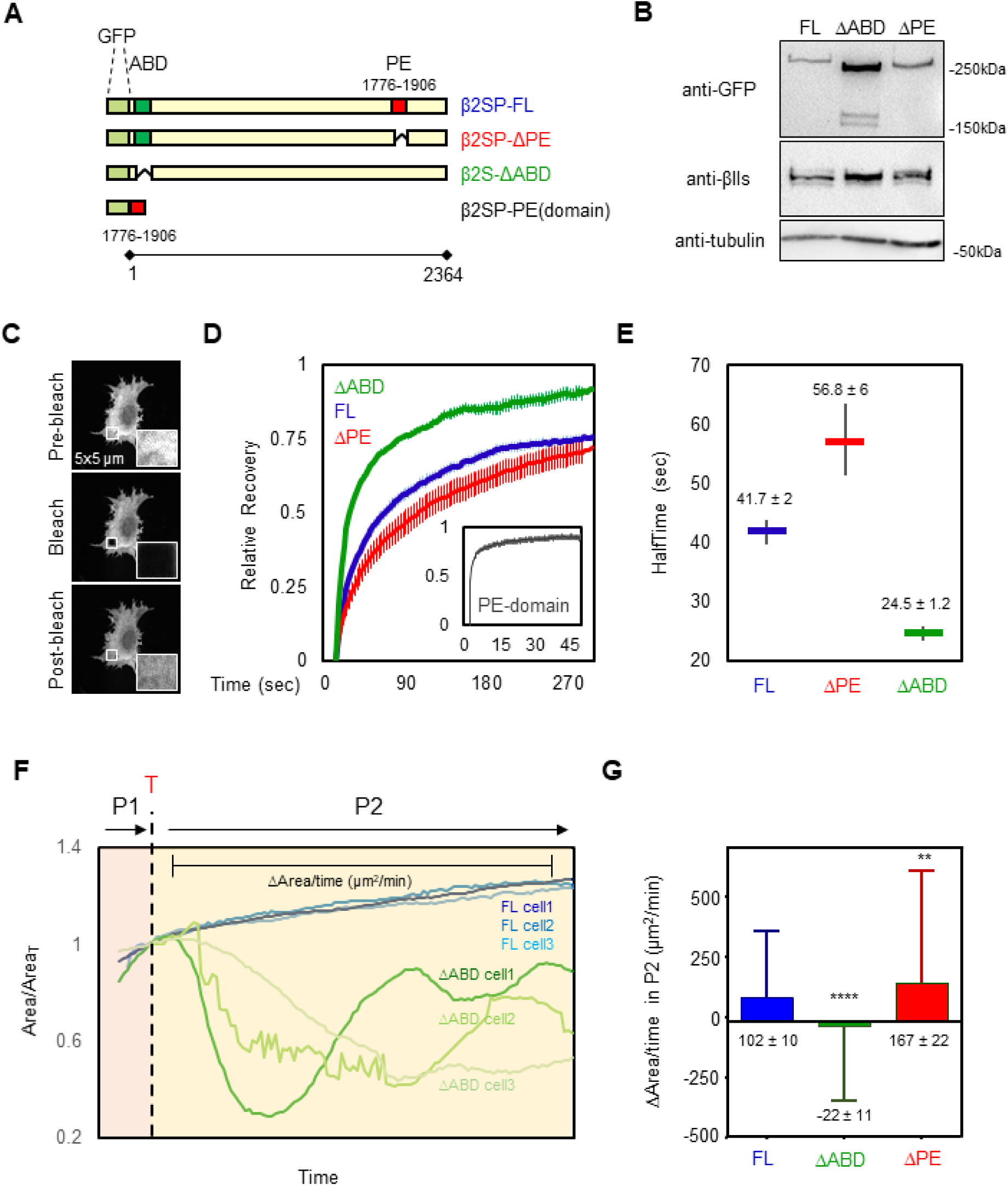
βII-Spectrin variants show different dynamics and properties during cell spreading. A) Cartoon representation of the βII-spectrin deletion mutants analyzed in this study. B) Total cell lysates of MEFs expressing exogenous GFP-βII-spectrin variants analyzed by anti-GFP and anti-βII-spectrin antibodies in western blot assay. C-D) FRAP assay of the βII-spectrin deletion mutants expressed in MEFs. The fit to a single exponential equation is shown (n= 15-20 cells, mean±SEM), and the resulting half time recoveries are plotted in E (mean±95% confidence interval). Full-length protein displays 74.8% of mobile fraction and half time recovery of 41.7 seconds, ΔABD 87.2% mobility and 24.5 seconds recovery and ΔPE 73.2% and 56.8 seconds. PE-domain only resulted in 87% mobility and 1.45 seconds recovery (inset). F) Normalized cell area growth during P2: three stereotypical MEFs transfected with GFP-βII-spectrin FL are plotted in blue, while MEFs expressing ΔABD are shown in green and followed for 10 minutes after P1/P2 transition (T) by live TIRFM. G) Quantification of ΔArea/time extracted from each frame of time-lapses during 10 minutes after transition into P2 (FL n=792 frames, ΔABD n=840 frames, ΔPE n= 502 frames, 7-9 cells. Mean±SD, one-way Anova statistical analysis with multiple comparison, **p<0.008 ****p<0.0001).

When looking at the cell shape remodeling mechanisms during spreading, ΔABD expressing cells underwent a normal P1 phase, while several collapses of protrusions were observed during the contractile phase (P2, Movie 4 and 5). In accordance with our FRAP results, spectrin meshwork cohesion through actin protofilaments binding appears instrumental to sustain PM when contractility is at play (Movie 4 and 5). These collapsing events were different from the retractions described earlier where spectrin replaced the actin-based support to the lipid bilayer. Here, simultaneous collapses of actin and spectrin were observed (Figure 5 F and Figure S4 E-F), followed by further attempts of the cell to spread over the substrate. As a consequence of these set of events, we recorded a negative Δarea/min rate since positive values were offset by negative events (during the 10 minutes spreading phase after the transition in P2, Figure 5 F-G). On the contrary, FL-βII-spectrin expressing cells displayed a stereotypical steady increase in area during P2; ΔPE expressing cells spread even faster than the FL-expressing cells and retraction zones highlighted with a remarkable ΔPE-spectrin accumulation (Figure S4, Movie 4).

Altogether these results indicate that the binding of βII-spectrin to actin protofilaments rather than PM is key for a correct meshwork dynamic during myosin II driven contractility, conferring resilience to the cell.

### Spectrin is condensed in actin-poor retracting zone under cell-stretch and is depleted in actin-rich blebs induced by compression

After establishing the dynamic response of spectrin during cell shape re-arrangement, we tested the reaction of the spectrin meshwork to perturbation of cell mechanics induced by stretching and compressive stresses. Indeed, if the spectrin meshwork condensation was a general mechanism to preserve cell and PM integrity (as shown in Figure S4 for cell shape changes), it may also display similar dynamic behavior under environmentally-driven mechanical perturbations. To test this hypothesis, MEFs were seeded on a deformable silicone membrane. Polygonal cells, characterized by the presence of long arcs between adhesive protrusions, were monitored during biaxial stretch on a custom-built device that could impose an increase in the stretch of up to 30% of the initial area (Figure 6 A-B, see methods). Since silicone membrane limits the possible imaging methods to wide-field illumination, we excluded moderate to high overexpressing cells from the analysis to avoid artefacts. Under stretching, GFP-βII-spectrin-expressing cells retained most of the prominent adhesions onto the substrate, while actin treadmilling activity at lamellipodia was blocked (visualized by Lifeact-RFP, Figure 6 B, double asterisks), as we previously reported (Pontes *et al*., 2017). Consistent with our previous observations, spectrin signal sharpened in the arc-shaped zones under progressive stretch, creating a frame around the cell that disappeared when the stretch was released (Figure 6 B).

**Figure 6.**
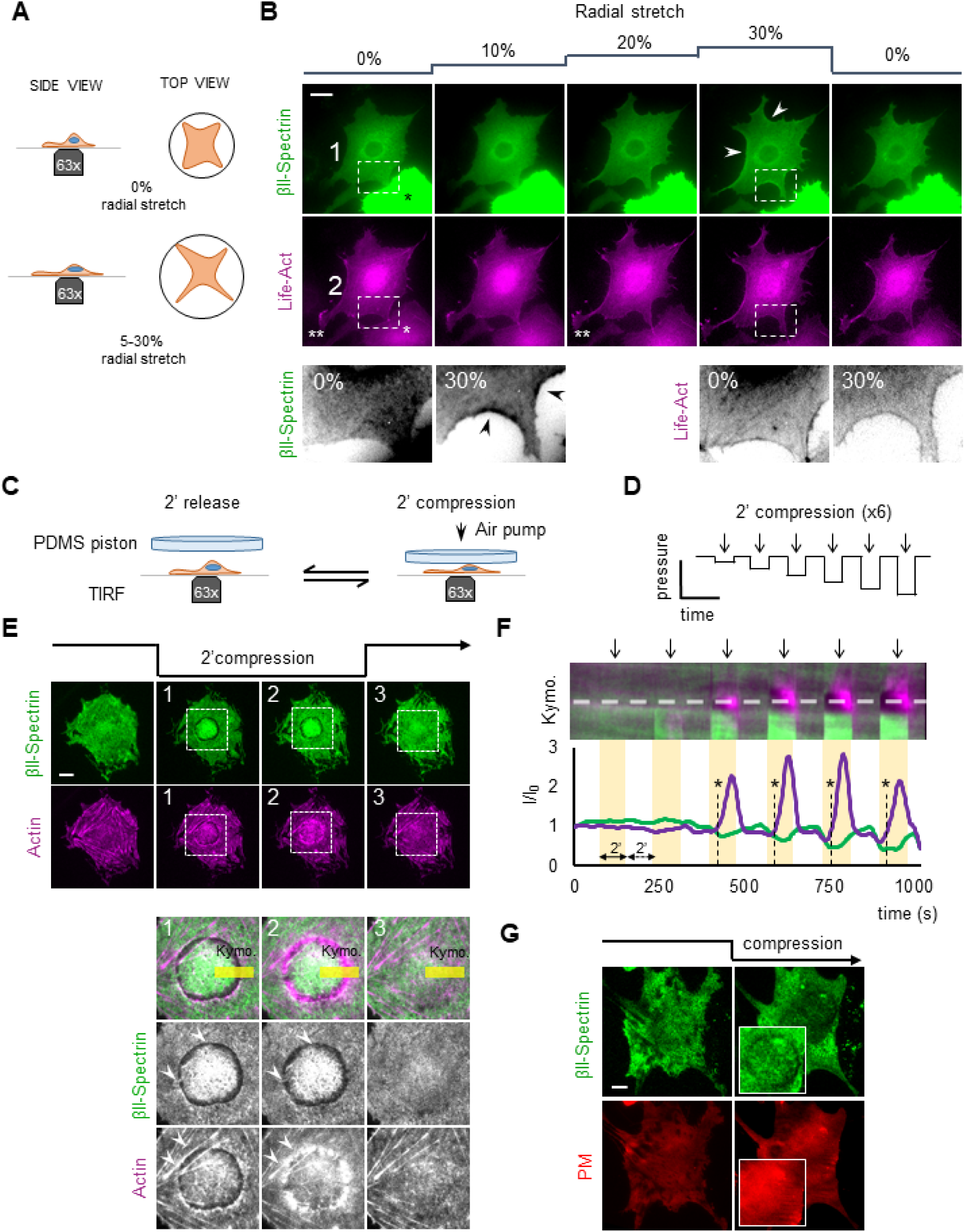
βII-Spectrin reactions to mechanical perturbations highlight the interplay with Actin. A) Cartoon representation of the cell stretching device implemented in this study. B) MEFs transfected with GFP-βII-spectrin (green) and LifeAct-RFP (magenta) seeded on the fibronectin-coated silicone membrane and stretched bi-axially (0-30%) during live EPI fluorescence imaging (white asterisk indicates transfected MEFs with high intensities excluded from the analysis, scale bar: 20 μm). Double white asterisks highlight a lamellipodia blocked during the stretching. In the dashed boxes 1-2, representative cell edge behavior observed among independent experiments, highlighting peculiar condensation of βII-spectrin in curvature zones not enriched by actin (arrowheads) at the maximal stretch (30%). C-D) Cell compression set-up and the applied step-increase protocol are schematized. E) GFP-βII-spectrin (green) and RFP-actin (magenta) expressing MEFs are imaged by live TIRFM during the entire compressive protocol. Four relevant time points are shown: pre-compression, early and late compression, and during the release phase (scale bar: 10 μm). Key details consistently observed between independent experiments are highlighted by dashed boxes and are zoomed in panel 1-2-3. The reaction in correspondence of the nuclear edge, brought into the TIRF plane by the compressive stress (white arrowheads), is quantified by the kymograph analysis (F) over the yellow rectangle. Fluorescence intensities across the dashed line in F are plotted in the graph; clearance of βII-spectrin and the delayed actin polymerization is observed (asterisks and dashed lines in the graph). G) Control cells expressing PM-marker (red) and βII-spectrin (green) are subjected to the same compression/relaxation protocol. Insets focused on the cortex underneath the nucleus, where PM-marker retains its continuity (scale bar: 10μm).

Further insights were also provided by the brutal detachment of less strongly attached adhesions in cells subjected to the gradual stretching protocol. Differently from the cell-driven retractions observed during spreading assay, actin and spectrin scaffolds condensed simultaneously and colocalized in the collapsed zone (Figure S5 F-H). Notably, this was the only mechanical events leading to apparent colocalization between the two cytoskeletons. We interpreted that in those particular fast events, actin and spectrin meshworks react passively as opposed to all the other mechanical perturbations where the crosstalk between the two leads to fine-tuned adaptation and reorganization. Spectrin meshwork active condensation in arc-shaped membrane retractions (i.e. spectrin-rich membrane curvatures) might be a process that operates in the absence of local actin under extrinsic (stretch) as well as intrinsic (spreading/polarization, Figure S4) mechanical challenges.

Next, we monitored spectrin meshwork dynamics under compressive stresses. We built a custom device to apply longitudinal, uniaxial compression/relaxation cycles on single cells, which were monitored in real-time by TIRFM (Figure 6 C-G, Figure S5 C-E, Movie 6). The increase in intracellular pressure caused by the compressive strain affected cell cortex integrity and induced the formation of blebs (Figure S5 C-D). Direct readout of the applied force on single cells is not possible in our setup due to variations in cell height, therefore the compressive piston was gradually lowered until cells showed blebbing. Compression-induced blebs (i-bleb) clearly displayed the flow of cytosolic actin directed into the newly formed blebs, while the majority of the spectrin signal was retained in the cell body (Figure S5 C-E). Upon the release of the piston, i-blebs were resorbed into actin-enriched tubular-like structures devoid of spectrin.

These results show that spectrin and actin skeletons display clearly distinct spatio-temporal dynamics depending on the nature of the mechanical challenge. Upon fast event, like cell detachment, both the actin and spectrin meshwork passively condense forming an undistinguishable “plug-like” structure as cell retract. Instead, during controlled retraction, spectrin can condense in actin-poor zones and that appear to support the PM. On the contrary, in fast-protruding zones non-actively driven by the actin cytoskeleton, like in mechanically-induced blebs, spectrin can be uncoupled from the PM, potentially preserving cell cohesion and cytosolic content, while actin flows into the bleb and progressively polymerizes into defined structures, as previously observed (Charras *et al*., 2006).

### Spectrin, actin and plasma membrane create a continuous but dynamic composite material

Surprisingly, we consistently observed that azimuthal uniaxial compression caused the nucleus to act as a dissipating additional piston on the cell cortex facing the coverslip (visualized by TIRFM), inducing most of spectrin and actin reactions to occur underneath this organelle in relatively flat MEFs. To shed further light on this phenomenon and better control the magnitude of the compression, we applied cycles of compressions/relaxations (2+2 minutes) at increasing strength (Figure 6 C-D, Movie 6). Piston contact with the cell roof marked the first compression step and did not affect basal spectrin signal. The piston was further lowered increasing progressively the stress on the cell. As compression increased, spectrin fluorescence increased right under the nucleus (Figure 6 E-F and Figure S5 B). Simultaneously, an unexpected spectrin- and actin-depleted rim formed in correspondence of the nuclear envelope (Figure 6 E). Remarkably, *de novo* actin-polymerization characterized by concentric inward flow specifically occurred in this bare PM region within the 2 minutes of compression (Figure 6 F and Movie 6). Careful examination showed spectrin tethered by few fibrous stretches across the rim (Movie 6). The release of the compressive stress blocked actin polymerization and was followed by a fast disappearance of the actin speckles (Figure 6 E, Movie 6). Upon relaxation, spectrin reacted completely differently than actin, since it immediately closed the rim without leaving a track of the tear in the meshwork, entangling and fencing the few remaining actin speckles (Figure 6 E-F and Movie 6). Occasionally, a similar behavior could also be observed by compression-relaxation of large cytosolic vesicles (Figure S5 A). Co-staining of PM with spectrin indicated that the membrane kept its integrity during the entire compressive stress (Figure 6 G).

These results represent a direct experimental demonstration of our previous observations on the dynamic cooperation between actin and spectrin in the cortex under mechanical challenges. Indeed, spectrin acts as an elastic continuous meshwork which can be stretched and depleted locally, thus working as a fence for the actin skeleton. On the other hand, bare PM is not a stable condition and the spectrin/actin cortex is constantly trying to occupy cytoskeletal-free space by covering it like a fluctuating elastic “veil-like” structure (spectrin) or polymerizing on it (actin).

Finally, to study more directly the spectrin elastic behavior in supporting the PM, osmotic shocks were applied to the cells as a third paradigm of environmentally-driven mechanical perturbation. These experiments aimed to simulate cycles of stretch-relaxation of the PM, while allowing us to monitor βII-spectrin reactions. Mean fluorescence intensity changes were simultaneously recorded for βII-spectrin and a fluorescent PM marker over the projected cell area. Spectrin fluorescence alone, registered by TIRFM, showed reduction during hypotonic shocks and increase during isotonic relaxation, however, the ratio between βII-spectrin/PM signals did not significantly shift from the initial ratio during several subsequent cycles (Figure S6 A-B). When soluble GFP was used as non-membranous control, consistent reduction in the GFP/PM ratio during hypotonic shocks could be recorded (Figure S6 C-D), while dual-tagged GFP-βII-spectrin-mCherry displayed a constant ratio (data not shown). Interestingly, ratiometric images failed to display homogeneous intensity throughout the entire cell, suggesting zonal enrichment or depletion of one of the two components. Local analysis during osmotic shocks displayed an initial reduction in the βII-spectrin/PM ratio that was compensated during later shocks, while a second region of the same cell matched the linear ratio shown over the entire cell projected area (Figure S6 E-F). Active lamellipodia during isotonic recovery behaved as expected, displaying reduced βII-spectrin/PM ratio compared to the adjacent cell body (Figure S6 G). Conversely, under hypotonic shock, increase in PM tension abruptly blocked lamellipodia activity (Figure S6 G) as previously reported (Gauthier, Masters and Sheetz, 2012; Kosmalska *et al*., 2015).

Altogether these results support the existence of local redistribution mechanisms of the spectrin mesh at meso-scale level. We concluded that βII-spectrin elastic support of the PM at whole-cell level is maintained by keeping constant the ratio between the two components, while it can locally and transiently drift to allow the occurrence of specific PM-linked events.

### Endocytic dynamic integration in the spectrin/actin/plasma membrane composite

Spectrin dynamics and the complementary interplay with actin pointed out the ability of the two meshwork to create PM microdomains that are consistent with the revised fluid-mosaic model of PM organization (Kusumi *et al*., 2012). Spectrin has been associated with PM organization, potentially positioning clathrin-mediated endocytosis (CME) events at cell-cell junctions (Jenkins, He and Bennett, 2015). We tested whether spectrin was also involved in this mechanism in fibroblasts by providing molecular details of the clathrin pit distribution and dynamics in fixed and live specimens. By immunostaining analysis of endogenous clathrin-heavy-chain (CHC), βII-spectrin and actin, we found that the three molecular components were not colocalized and appeared rather mutually excluding each other (Figure 7 A, Figure S7 H). To strengthen this observation, multiple discrete clathrin structures were selected, registered in 2×2µm ROIs and clustered into two groups of different size (<300nm^2^ and 300-500nm^2^) following a recently published approach (Mund *et al*., 2018). The density maps displayed clathrin pits positioned at the center of spectrin-depleted zones surrounded by spectrin-rich areas. Remarkably, the diameter of the averaged spectrin-depleted zones almost matched in size the clathrin pits projections (Figure 7 B-C). While most of the high-intensity actin structures, such as stress fibers, were clearly distinct from the pits, the averaging of >100 pits led to the identification of a discrete actin enrichment in correspondence of the clathrin staining (Figure 7 A-C). These observations are fully consistent with the current maturation models of endocytic structures (Kaksonen and Roux, 2018; Kirchhausen, 2009). Our analysis indicates that a potential hindrance mechanism might be at work. According to this, the spectrin meshwork is able to delimit the zones of the assembly of clathrin pits.

**Figure 7.**
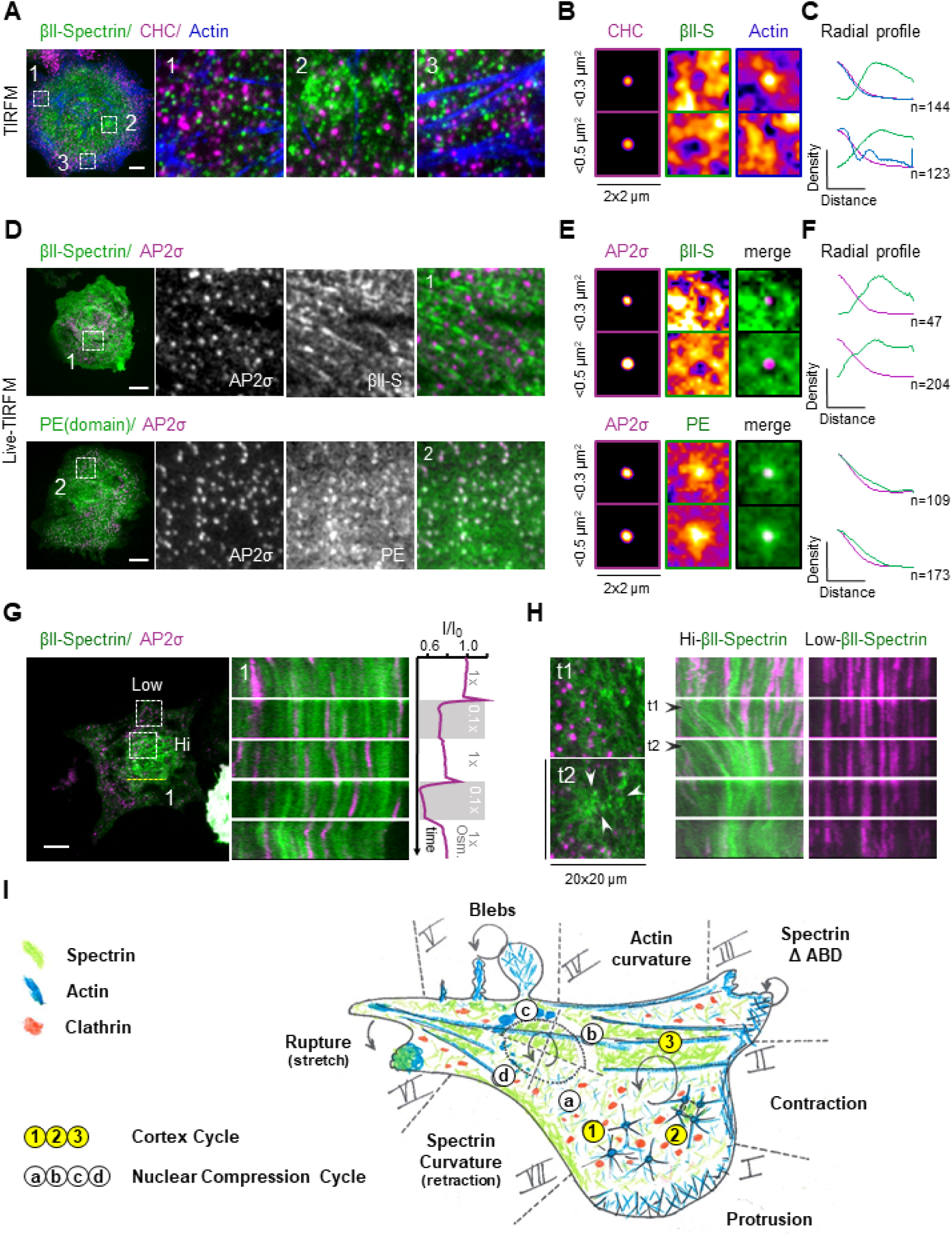
Clathrin Endocytosis dynamic integration in the spectrin/actin/plasma membrane composite. A) TIRF microscopy images of MEFs immunostained for endogenous βII-spectrin (green), clathrin heavy chain (CHC, magenta) and F-actin (blue, scale bar: 10 μm). Three different subcellular regions are highlighted (dashed boxes 1-3): high-density CHC zone (1), βII-spectrin-rich zone (2) and actin stress fibers 2 (3). B) Density maps generated by aligning discrete clathrin pits of small (<0.3 µm^2^ n=144) and larger size 2 (0.3-0.5 µm^2^, n=123) are shown, while in C the correspondent radial density profiles for the three channels are presented. D) Representative live TIRF microscopy images of MEFs transfected with GFP-βII-spectrin (green) and the clathrin adapter AP2σ-mCherry (magenta) (scale bar: 10 μm), Unsharp and Gaussian filters were applied. PE-only domain transfected fibroblasts show homogeneous PM localization and enrichment in correspondence of AP2σ pits, as shown by density maps (E) and radial profile analysis (F) generated as 2 2 2 previously described (GFP-βII-spectrin: <0.3 µm^2^, n=47 and 0.3-0.5 µm^2^, n=204; PE-domain: <0.3 µm^2^, 2 n=109 and 0.3-0.5 µm^2^, n=173. G) Representative time-lapse TIRFM images during osmotic shocks: kymograph is generated in correspondence of the dashed yellow line (1), where spectrin mesh and AP2σ pits display coordinated lateral motion in response to osmolarity changes. Whole-cell AP2 intensity signal in response to the osmotic shocks is plotted in the graph (vertical plot). H) 20×20µm ROIs are drawn at low- and high-spectrin density zones around AP2 pits (correspondent to dashed boxes in G, only zooms of Hi-density zone are reported at two different frames). Kymographs display the differential behavior of the pits observed during spectrin remodeling. (I) Model resuming our findings on the dynamic response of spectrin during mechanoresponse. (I) to (VII) highlight the cell edge behavior. (I) In cell protrusions actin polymerization dominates. (II) During contraction, actin is condensed and forms transverse arcs. In I and II spectrin is secluded and “passively” follow acto-myosin lead. (III) Deletion of the Actin binding domain of spectrin induces edge instability upon contractility activation. (IV) Mature actin bundles sustain the PM, spectrin is not recruited to those actin curvatures. (V) Upon cell compression blebs enriched with actin but devoid of spectrin are formed. While actin polymerizes and condense in the bleb, spectrin localizes and marks the former position of the cell edge before compression. (VI) Abrupt cell detachment induces a “plug-like” formation in which actin and spectrin seem to colocalize. (VII) In actin depleted but spectrin-rich edge curvature, spectrin condenses as the edge move inward, potentially holding the PM and responding to the increased membrane load. (a) to (d) highlight the cell cortex behavior under compressive stress. (a) to (b) The native acto-spectrin cortex gets cleared under the edge of the nuclear envelope upon compression. (c) During compression, in this gap of the cleared membrane, actin polymerization occurs and covers progressively the bare bilayer. (d) Upon relaxation, spectrin meshwork elastically recoils, entangling the polymerizing actin and restoring the original cortex organization. Stress fibers under the nucleus are not affected by the compression. (1) to (3) highlight the non-perturbed cell cortex behavior. (1) In spectrin-less zones, actin nodes can form. (2) These aster-like structures move and coalesce upon myosin II-mediated contractility. This mechanism synchronously modifies the local density of the spectrin meshwork, with expansion and condensation between coalescing nodes. Further acto-myosin condensation lead to bundles formation interleaved by spectrin-rich territories. (3) These high-density zones impede clathrin-mediated endocytosis (red dots), which otherwise occurs between spectrin fences. Those steps occur cyclically upon cell control or drugs/mechanical stimuli.

Live imaging analysis confirmed the exclusion of GFP-βII-spectrin from clathrin structures visualized by the adapter mCherry-AP2σ. Specifically, AP2-decorated pits appeared in the void patches that characterized the spectrin meshwork (Figure 7 D-E-F). Notably, the membrane-bound PE-domain of spectrin used here as negative control did not display the same behavior. CME is a highly dynamic and heterogeneous process with several layers of regulation, including membrane tension modulation by osmolarity (Boulant *et al*., 2011). Therefore, we monitored this process with respect to the dynamic spectrin re-organization during osmotic changes. As expected, decreased spectrin signal intensity during hypotonic shocks was accompanied by a fast-transient increase and followed subsequently by severely reduced AP2 intensity, which was restored after the transition to isotonic conditions (Figure 7 G, Movie 7). More interestingly, kymograph analysis revealed coordinated flow between the two channels during cell adaptation motion, suggesting that AP2 pits are hooked to the spectrin meshwork. The same effect was not observed for AP2 and PE-domain only (data not shown). We compared discrete AP2 pits in cellular zones characterized by high and low GFP-βII-spectrin densities during osmotic changes. Interestingly, when the spectrin cytoskeleton reorganized into large condensation zones, several pits disappeared from the TIRF plane, most likely engulfed by the fencing capability of the spectrin meshwork (Figure 7 H, Movie 7). This was not observed in low spectrin-density zones of the same cell, indicating that a local hindrance mechanism might be operative.

Altogether these results support a critical involvement of spectrin spatio-temporal reorganization in the positioning of endocytic structures.

## Discussion

Here, we provide a universal view on how ubiquitous and evolutionary conserved spectrin dynamically interplays with acto-myosin, the lipid bilayer and the endocytic machinery to sustain the PM during intrinsic and extrinsic mechanoadaptative events. We recapitulated our main findings in the working model in Figure 7 I.

Our analysis of a variety of mammalian cells growing under various geometrical constraints, suggests that there are discrete PM territories supported either by an actin scaffold or by a spectrin skeleton.

Dynamic studies in fibroblasts spreading onto adhesive substrate unveiled the assembly mechanism governing spectrin meshwork organization during the early phase of cell-substrate interaction. Upon activation of acto-myosin contractility, coordinated motion of spectrin and coalescent actin nodes emphasize the interplay between the two scaffolds during full maturation of the cytoskeleton (Figure 7 I, cortex cycle 1 to 3). Remarkably, we observed spectrin meshwork dynamics to rely on myosin II contractility either during cytoskeletal maturation as well as in established cell cortex. Current models describing the transmission of myosin-dependent contractility at isotropic cortex hardly explain how forces exerted on non-polarized scaffolds can produce homogenous movement. Since cortical actin protofilaments might be too short and rigid to generate coherent contractility (Koenderink and Paluch, 2018), the hierarchical actin/spectrin organization and the cohesiveness provided by the described meshwork have the potential to reconcile this paradox. This is highlighted by the expression of the spectrin ΔABD mutant, documenting edge instability characterized by synchronous spectrin/actin retraction (Figure 7 I, zone III). We speculate that in the actin-poor but spectrin-rich lamella, the spectrin meshwork can act as a force-transmitting “veil-like” structure underneath the PM. This veil creates a continuum at the lamellipodia/lamella border with the contractile structure localized deeper in the cell body. The dominant-negative ΔABD expression might, thus, uncouple these distinct frameworks and create a mechanical discontinuity in this cohesive architecture. However, also long-term perturbation of the other cytoskeletal systems, such as microtubules, can affect the organization and the cohesiveness of the spectrin meshwork and associated-proteins (Jenkins, He and Bennett, 2015), albeit with different mechanisms and time-scale that require further investigations.

Our results during extrinsic mechanical perturbations suggest that spectrin works as a sum larger than its individual parts (dimers and tetramers) and reacts differently depending on the nature of the applied mechanical cues. We provide further support to a fencing mechanism brought about by macromolecular condensation upon mechanical stimuli, firstly proposed in neuronal axons under compressive and bending forces, rather than a molecular stretch/relaxation model based on intramolecular distance discrepancies between different EM and super-resolution studies, (Krieg, Dunn and Goodman, 2014; Krieg *et al*., 2017).

Altogether, our results consistently highlighted the opposite polarity between the spectrin skeleton and actin, either during extrinsic perturbation or intrinsic cell polarization. To the best of our knowledge, this is the first dynamic and mechanistic description of actin/spectrin dualism during cell shape remodeling events. Based on different mechanical perturbations of the PM, we propose that spectrin is maintained globally at a constant ratio with the lipid bilayer. Locally, it reacts instead to intrinsic cues, such as PM collapse (Figure 7 I, zone VII), or to external perturbations, such as cell-cell fusion (Duan *et al*., 2018). These reactions correspond to a meshwork condensation rather than de-novo recruitment, as a self- and cell-protective mechanism. The actin/spectrin coordinated dynamics is particularly exploited during spectrin cortex clearance induced by mechanical compression of the cell, which triggered an actin nucleation-dependent protective mechanism in response to spectrin displacement (Figure 7 I). Together with spectrin exclusion from protrusive lamellipodia (Figure 7 I, zone I), these results suggest the existence of a previously neglected interference mechanism that hinder actin polymerization in presence of a spectrin-enriched cortex.

Our observations during different environmental perturbations strongly support the existence of a non-Brownian diffusion mode of the spectrin skeleton through the PM (Frick, Schmidt and Nichols, 2007). Indeed, the spectrin meshwork defines PM microdomains able to constantly remodel in response to external and internal cues. Such capacity integrates well into the revised “three-tiered meso-scale” version of the fluid-mosaic model of PM organization (Nicolson, 2014; Kusumi *et al*., 2012) and can complement the so-called “picket and fencing” mechanism prominently led by the actin cortex. Here, we implemented this model by adding the spectrin-rich territories in the context of membrane dynamics and topological microdomain organization. As previously hypothesized from biochemical data (Jenkins, He and Bennett, 2015), we observed membrane trafficking events, such as CME, taking place at PM microdomains “hamstrung” by spectrin, while pits maturation sustained by actin polymerization occurred specifically within spectrin fenestration (figure 7 I, cortex cycle 1 to 3). Of note, several mechanisms have been identified in the regulation of CME, many of them showing discrepancies and controversy with one another (Doherty and McMahon, 2009). Thus far, none of them has clearly taken into consideration the role of the spectrin meshwork and its local re-organization. Such a role in organizing PM trafficking events is consistent with a recent report proposing spectrin as a general ruler for cell-cell adhesion molecules organization in neurons (Hauser *et al*., 2018). Since it is reasonable to imagine similar mechanisms for membrane trafficking pathways by opposite directionality, we propose that the highly dynamic composite nature of the cortex under mechanoresponses is mainly regulated by an orchestrated “*menage a 4*” between PM, spectrin, exo/endocytosis and acto-myosin contraction-polymerization (see our conclusion model Figure 7 I). More generally, these results indicate that the spectrin skeleton dynamics is critical to shape and coordinate many PM-linked cellular processes in physiology and pathology.

## Acknowledgements

We are grateful to the IFOM imaging facility personnel, in particular D. Parazzoli, M. Garrè and E. Martini for technical support. IFOM cell culture facility and qPCR facility personnel. We thank Jagadish Sankaran (NUS) and Toh Kee Chua (NUS) for the preliminary experiments of this project. We thank MBI protein core facility personnel, in particular Chen Hongying for providing the constructs. We thank Paulina Nastaly (IFOM) for the help with micropatterning. We thank Mike Sheetz (MBI), Marco Foiani (IFOM), Sarah Barger (NYU) and all the members of Gauthier’s, Scita’s and Maiuri’s groups for helpful discussion. We thank Sara Sigismund (IEO) for kindly providing additional constructs. This work was supported by IFOM starting package, Mechanobiology Institute of Singapore grant WBS R-714-016-007-271, and the Italian Association for Cancer Research (AIRC), Investigator Grant (IG) 20716 to NCG, by H2020-MSCA individual fellowship to AG (796547) and by Fondazione Umberto Veronesi (FUV) doctoral fellowship for CG.

## Author contribution

AG, NCG contributed to the conception and design of the experiments, interpretation of data, drafting and critical revision of the article. AG, CG, NCG, QL, FA, PM performed the experiments. QL and FA conceived the engineered devices and helped with the experiments. AG, NCG, MAF, PM contributed to data analysis. All authors critically revised and approved the last version of the article.

## Extended data

**Movie 1** Fibroblast spreading assay: cell edge analysis of GFP-βII-spectrin and RFP-Actin

**Movie 2** Fibroblast spreading assay: PIV analysis of GFP-βII-spectrin and RFP-Actin flows

**Movie 3** Actin and βII-spectrin dynamics during Latrunculin A and Blebbistatin washout experiments

**Movie 4** Differential βII-spectrin deletion mutants’ behavior during spreading

**Movie 5** βII-spectrin-ΔABD displays edge instability during spreading

**Movie 6** Cell compression assay

**Movie 7** Meso-scale dynamic of GFP-βII-spectrin and mCherry-AP2 during osmotic shocks

**Figure S1.**
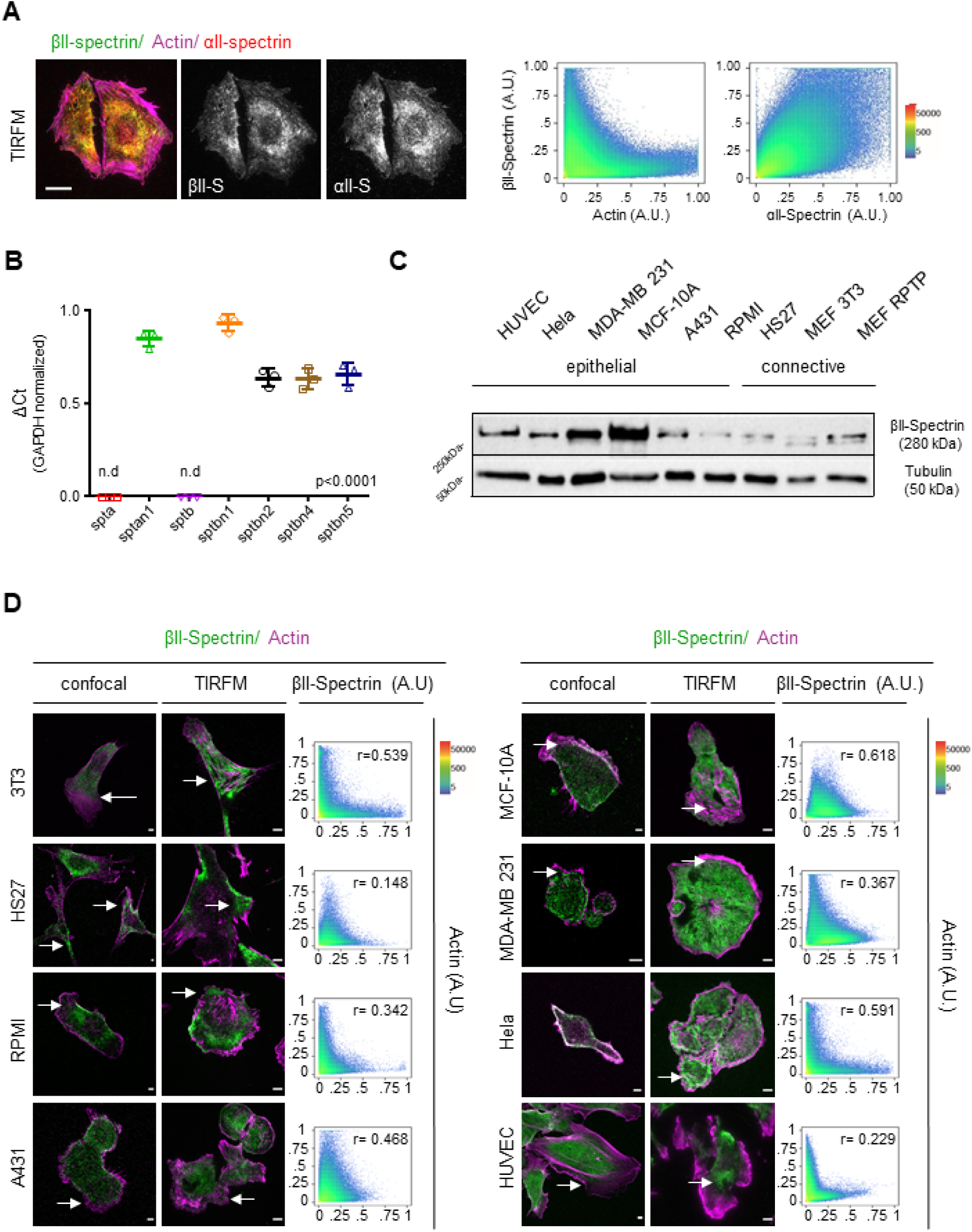
βII-Spectrin/Actin complementarity is observed in multiple cell lines by different backgrounds. A) Representative images of MEFs immunostained for βII-spectrin (green), αII-spectrin (red) and F-actin (magenta), visualized by TIRFM (scale bar: 10 µm). Correlation analysis of pixel intensities between the different channels is reported in the two scatter plots. B) Gene expression analysis of different spectrin isoforms in MEFs used for this study, measured by qPCR (n=3, n.d.=not detected, Statistical analysis: one-way ANOVA with multiple comparisons, p < 0.0001). The genes *sptan1* and *sptbn1* encode for αII and βII spectrin respectively. C) βII-spectrin protein expression in different cell lines analyzed by western blot in total cell lysates. D) Different cell lines were immunostained for F-actin (magenta) and endogenous βII-spectrin (green). Confocal and TIRFM images are presented, arrows indicate peculiar complementary zones (scale bar: 10 µm). Cross-correlation analysis by scatter plot between the two channels and Pearson’s correlation coefficients are reported for TIRFM images.

**Figure S2.**
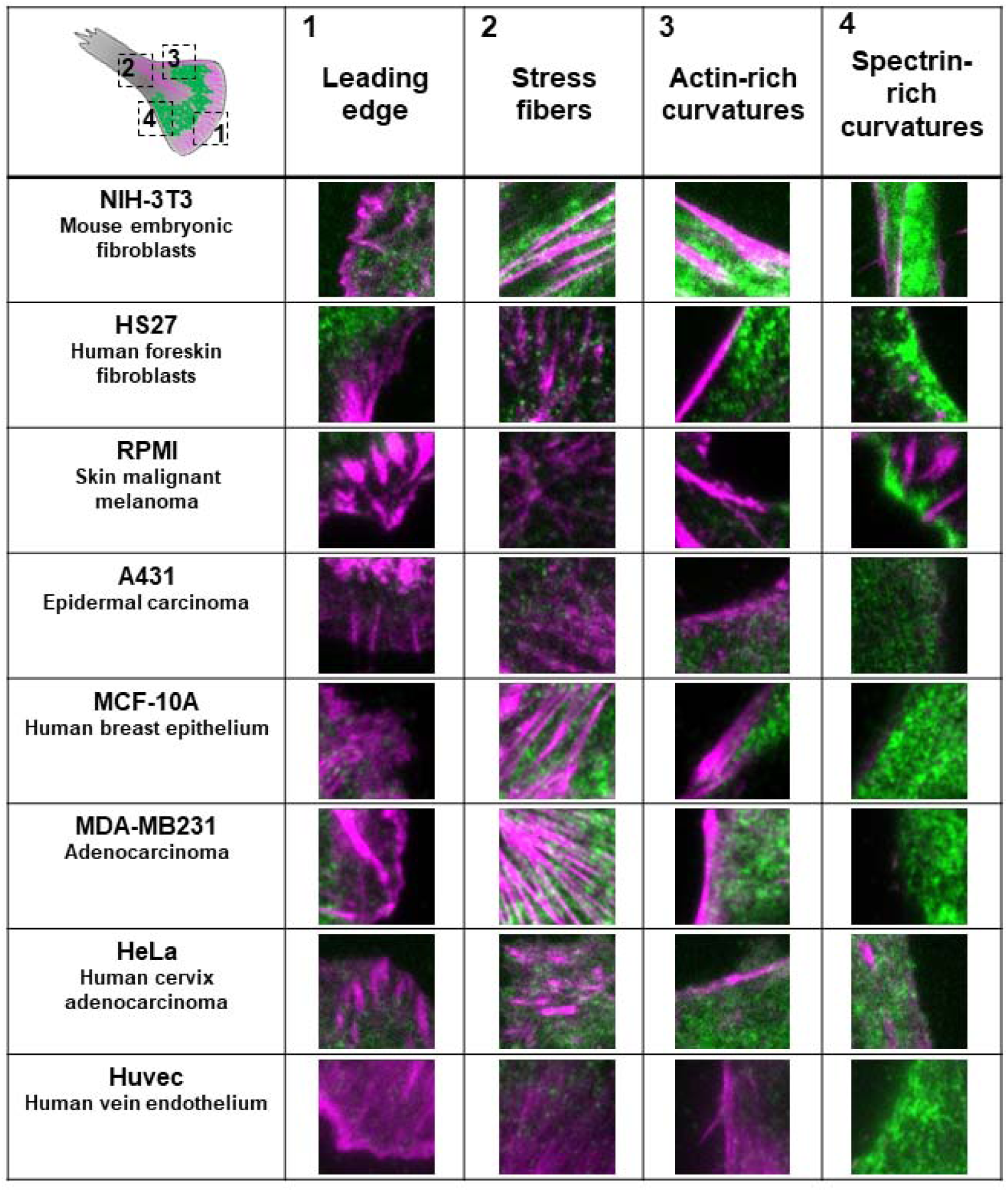
Zooms of βII-Spectrin and Actin complementary territories. TIRFM images (2×2 μm) of the different cell lines immunostained for endogenous βII-spectrin (green) and F-actin (magenta). Cell zones (as listed in Figure 1 A for MEFs) highlighting βII-spectrin/actin complementarit are reported.

**Figure S3.**
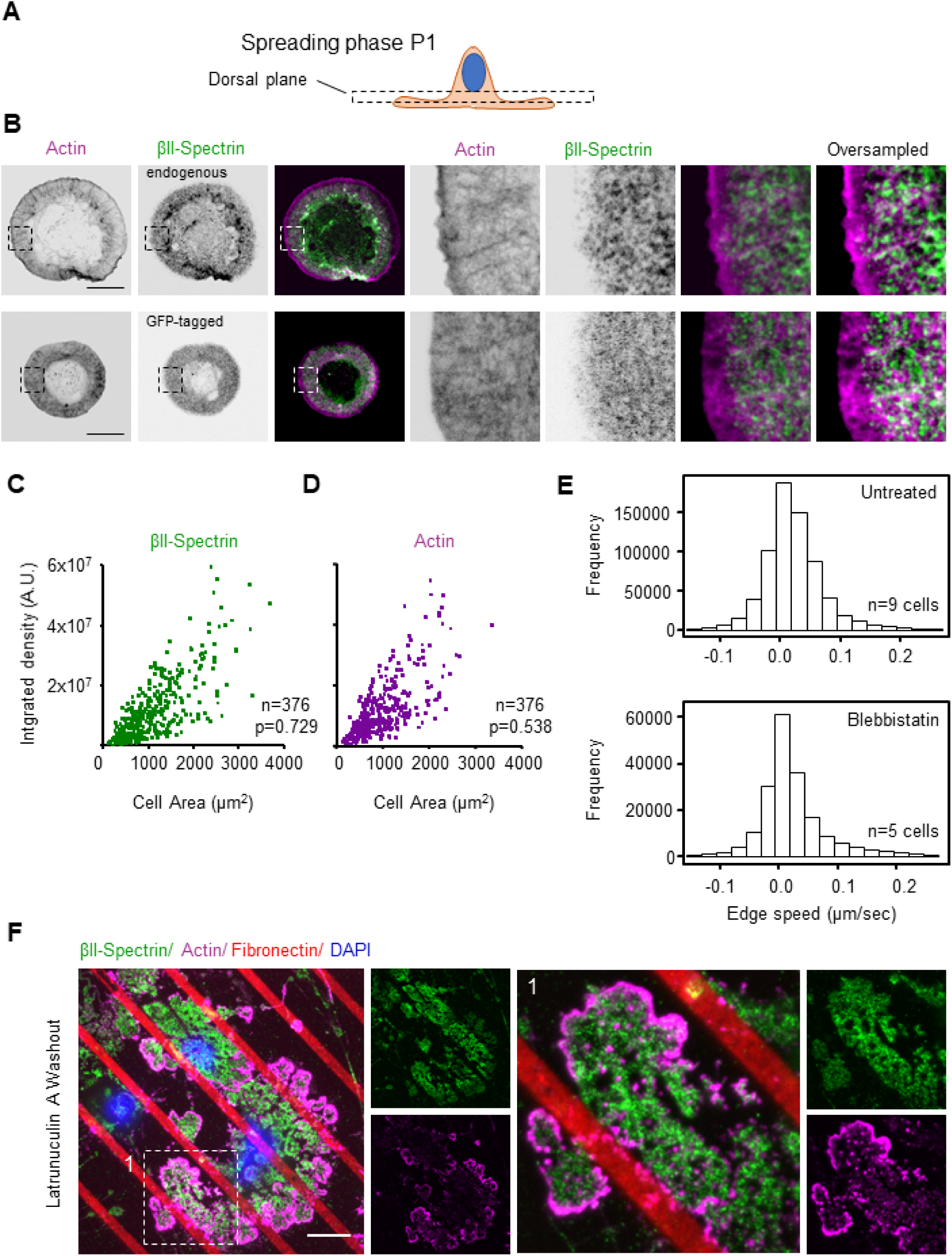
βII-Spectrin organization in P1; βII-Spectrin/area relationship during spreading; βII-Spectri and Actin recovery after LatA washout in cortex-mimicry zones. A-B) MEFs fixed during P1, immunolabelled for endogenous F-actin (magenta) and βII-spectrin (green, endogenous and GFP-tagged), and analyzed by 3D confocal microscopy. Optical sectioning is optimized to resolve the cortex on the cell dorsal plane during P1, as shown in the cartoon (scale bar: 20 µm). C-D) Cells fixed at different time points after seeding (between 5-20 minutes) and immunolabelled for endogenous βII-spectrin and F-actin. Projected cell area and fluorescence integrated intensities in TIRFM for the two proteins are reported, displaying linear correlation (n=376 cells, see Extended Table 1). E-F) Total data point distribution of the graphs displayed in Figure 2E and 2G, outliers were excluded from the analysis (threshold 0.0007; untreated: n=9 cells; blebbistatin: n=5 cells). Both analyses showed Gaussian normal distribution between the physiological speed range of −0.1 µm/sec and +200 µm/sec. F) MEFs seeded on micropatterned fibronectin-coated lines. TIRFM images of cells fixed during the washout phase after Latrunculin A treatment are shown (green: βII-spectrin, magenta: F-actin, red: fibronectin. Scale bar: 20 µm). DAPI (blue) is visualized in EPI mode to discriminate intact cells from debris. The white dashed box (1) is zoomed to highlight peculiar actin nodes formation in the non-adhesive cell cortex.

**Figure S4.**
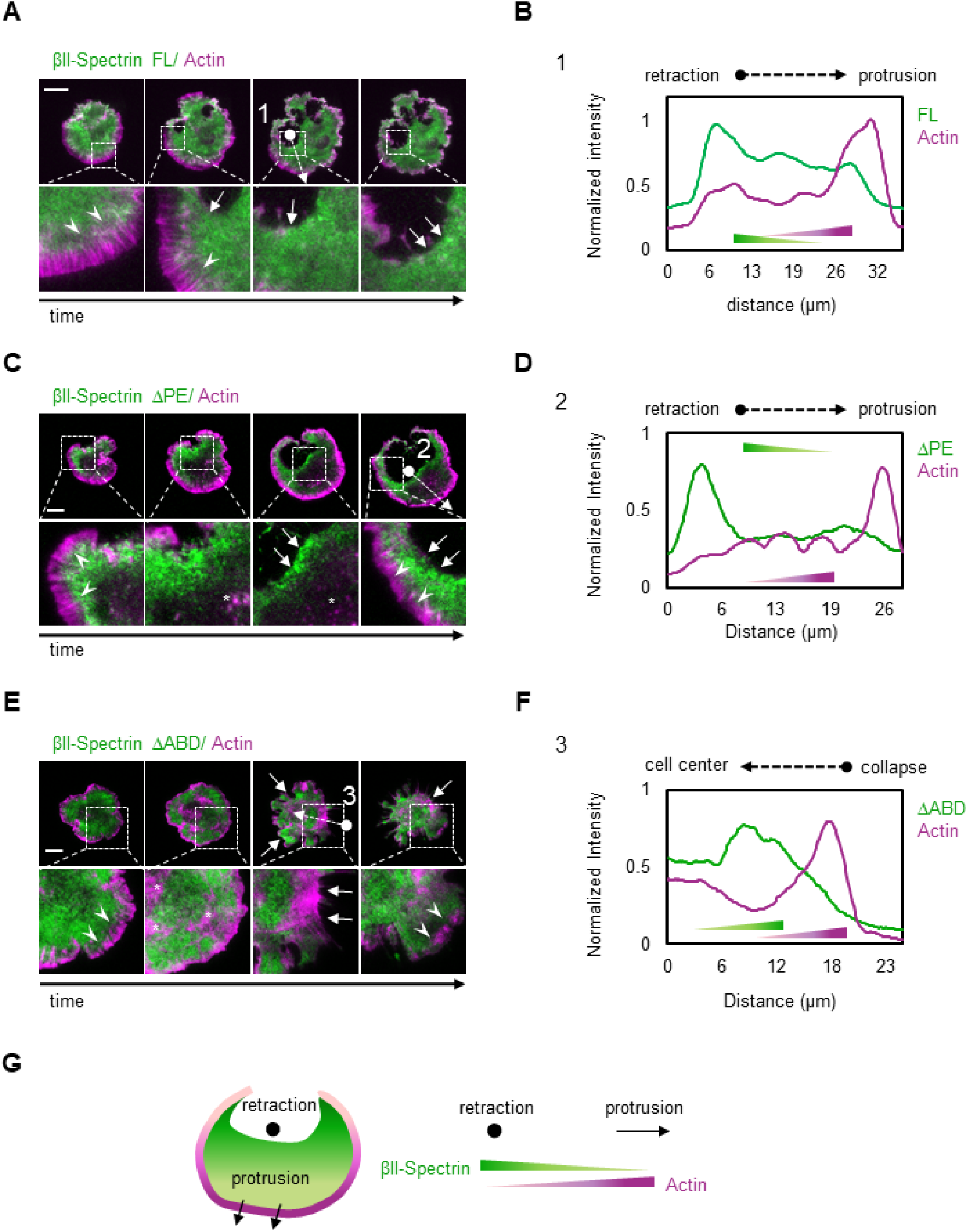
βII-Spectrin FL, ΔPE and ΔABD accumulate at “cracking” zones that spontaneously form durin cell spreading/polarization. A-C -E) Representative images of spontaneous retractile events observed in MEFs expressing GFP-βII spectrin variants during cell spreading by live TIRFM (GFP-βII-spectrin variants in green and RFP-Actin in magenta, scale bar: 20 μm). Relevant events are highlighted by the dashed boxes and zoomed in the lower panels: protruding zones are indicated by white arrowheads, retracting zones by white arrows. Line scan analysis of arrows with circular ends (1-2-3) are reported in B-D-F for both proteins, directionality indicated by black arrows. G) Cartoon model of Actin/βII-spectrin opposite polarity during protrusion/retraction events during the polarization phase of cell spreading.

**Figure S5.**
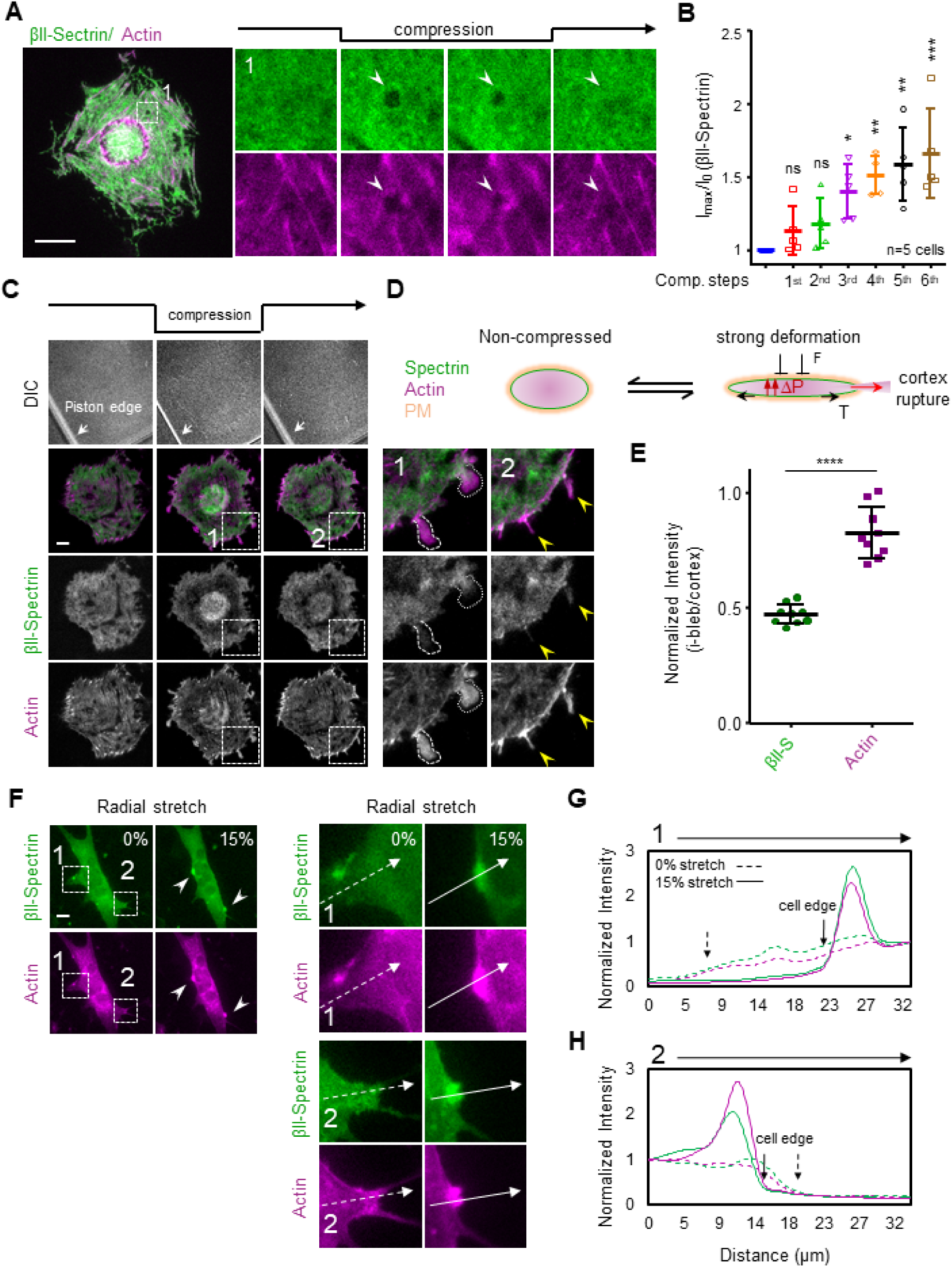
βII-Spectrin reactions to mechanical perturbations highlight the interplay with Actin also in vesicles, blebs and collapsed protrusions. A) The same cell presented in Figure 6 E is reported (scale bar: 20 μm). Zoom 1 highlights the appearance of a vesicular structure (white arrowhead) upon compression, devoid of both βII-spectrin and actin, resembling the clearance effect observed underneath the nucleus. B) Quantification of GFP-βII-spectrin maximal intensity at the cell body during the sequential compression protocol, normalized to the pre-compression phase (n=5 cells, mean±SD, one-way Anova with multiple comparisons * p<0.05, ** p<0.005, *** p<0.0005). C) Maximal compression experiments of GFP-βII-spectrin (green) and RFP-actin (magenta) expressing MEFs: compression stress is gradually increased until bleb formation is induced. Model of cortex rupture mechanism is schematized in D: key elements are the variation in intracellular pressure (ΔP) and cortex tension (T) during compression. Representative images are shown during (1) and upon release of compression (2), when induced blebs are resorbed into tubular-like actin enriched structures (yellow arrowheads). Actin and βII-spectrin content in the blebs compared to the adjacent cell body are quantified in E (n=9, mean±SD in 3 independent experiments, unpaired T-test: * p<0.0001). F) Bi-axial cell stretching experiment of GFP-βII-spectrin (green) and RFP-actin (magenta) expressing MEFs, imaged by live EPI fluorescence microscopy. Dashed boxes 1 and 2 highlights two specific cell protrusions that detached upon 15% stretch. Intensity profile across these processes (white dashed (0%) and full (15%) arrow) are plotted in G and H.

**Figure S6.**
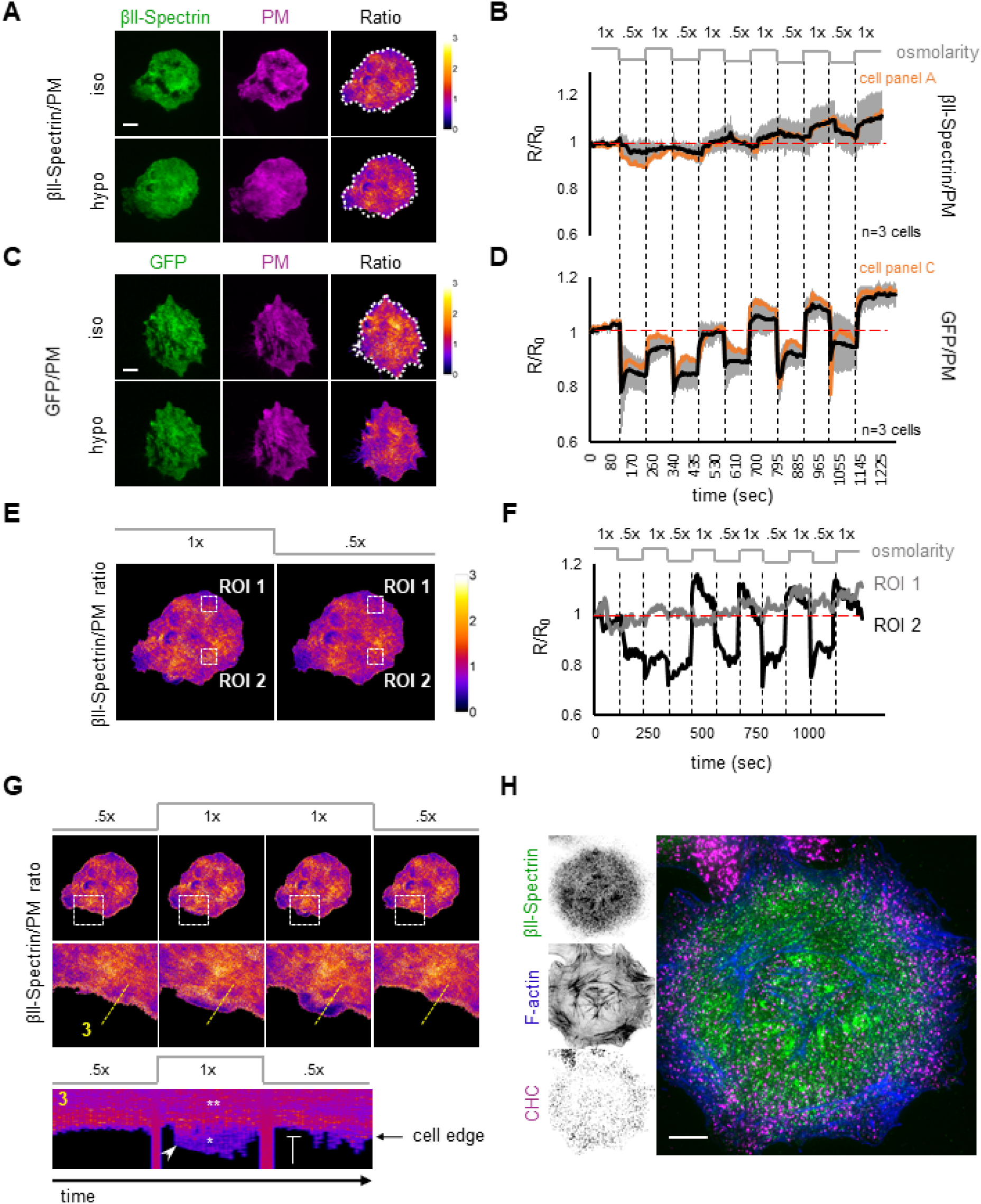
βII-Spectrin reactions to osmotic changes: global versus local behavior. A) Representative images of GFP-βII-spectrin (green) and PM-marker (magenta) transfected MEFs observed by live TIRFM during osmotic shocks. Five isotonic (1x)-to-hypotonic (0.5x) cycles were applied (B). Initial fluorescent signals are normalized to obtain the non-stoichiometric ratio βII-spectrin/PM (LUT fire, scale bar: 10 μm). The average ratio is plotted in B (black line, n=3 cells, mean±SD), while the ratio of the cell in A is shown by the orange line. C-D) As a positive control, the same protocol is applied to MEFs transfected with soluble GFP (green) and PM-marker (magenta), while GFP/PM ratio is plotted in D (n=3 cells, mean±SD). Zonal Ratio analysis at two extreme cases is reported in E: ROI1 presents high βII-spectrin/PM ratio, while ROI2 displays a lower ratio. As shown in the graph (F), the two ROIs behave differently: while ROI1 reacts similarly to the whole-cell analysis graph presented in B, ROI2 shows an initial decrease of the ratio sustained during the first two iso-to-hypotonic cycles, followed by a compensatory effect that restored the initial ratio during the last four cycles. A similar effect in a different PM zone is presented in G: lamellipodia (dashed box and zoomed in the bottom panel) characterized by high actin and low spectrin content are blocked during hypotonic shocks. Kymograph generated across the dashed yellow line (3): βII-spectrin/PM ratio is low in lamellipodia (* asterisk) compared to the adjacent cell body (** asterisks), and lamellipodia blockage during the hypotonic shock is observed. H) The same cell presented in Figure 7 A is shown: endogenous βII-spectrin (green), clathrin heavy chain (CHC, magenta) and F-actin (blue) and imaged by TIRFM (scale bar: 10 µm).

**Figure S7.**
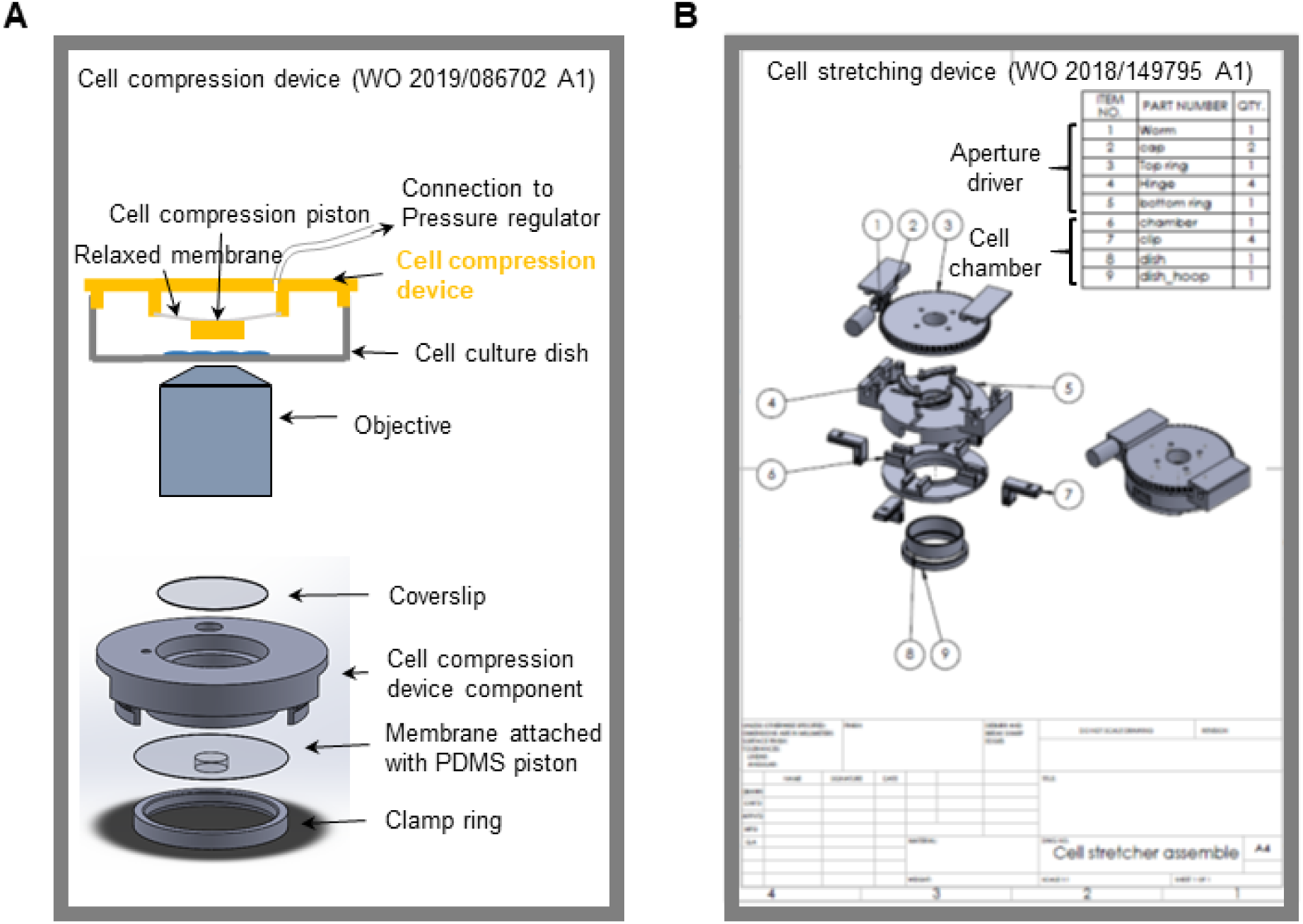
Cell compression and stretching devices. A) Schematic explosion illustrating the single components of the pneumatic Cell Compression device. The same is reported for the Cell Stretching device implemented in this study (B).

## Extended Data Tables

**Table 1.** Spreading area/spectrin and area/actin linear regression analysis.

|  | Actin Integrated density | Spectrin integrated density |
| --- | --- | --- |
| Pearson correlation coefficient | 0.5386254 | 0.7296664 |
| Linear regression |  |  |
| Best-fit values |  |  |
| B0 | 4.758e+02 | 4.708e+02 |
| B1 | 4.459e-05 | 4.405e-05 |
| Std. Error |  |  |
| B0 | 4.260e+01 | 2.821e+01 |
| B1 | 3.962e-06 | 2.345e-06 |
| T value |  |  |
| B0 | 11.17 | 16.69 |
| B1 | 11.26 | 18.79 |
| Goodness of Fit |  |  |
| Degrees of Freedom | 371 | 301 |
| Multiple R-squared | 0.2901 | 0.5324 |
| Adjusted R-squared | 0.2878 | 0.5309 |
| p-value | 2.2e-16 | 2.2e-16 |
| Beta estimate | 4.45906e-05 | 4.40486e-05 |
| Standard error | 3.96159e-06 | 2.344543e-06 |
| T value | 11.25574 | 18.78771 |
| P value | 7.35929e-25 | 4.67908e-53 |
| F statistic | 7.16465e-25 | 7.16465e-25 |
| F | 4.45906e-05 | 4.40486e-05 |
| Model p | 3.96159e-06 | 2.34454e-06 |
| Outliers | 12 | 12 |

**Table 2.** Dual FRAP Analysis.

|  | Diamide |  | Nocodazole |  | Latrunculin A |  | Blebbistatin |  |
| --- | --- | --- | --- | --- | --- | --- | --- | --- |
|  | Control | Treated (5') | Control | Treated (5') | Control | Treated (5') | Control | Treated (5') |
| One-phase association |  |  |  |  |  |  |  |  |
| Best-fit values |  |  |  |  |  |  |  |  |
| Y0 | 0.009453 | 0.01164 | -0.05037 | 0.02238 | -0.01107 | -0.08307 | -0.004542 | -0.002546 |
| Plateau | 0.8569 | 0.3446 | 0.8513 | 0.8262 | 0.8294 | 0.8526 | 0.8371 | 0.6676 |
| K | 0.01803 | 0.01437 | 0.0196 | 0.01504 | 0.01989 | 0.02544 | 0.02008 | 0.01698 |
| Tau | 55.45 | 69.59 | 51.01 | 66.48 | 50.29 | 39.3 | 49.79 | 58.91 |
| Half-time (sec) | 38.44 | 48.24 | 35.36 | 46.08 | 34.86 | 27.24 | 34.51 | 40.83 |
| Span | 0.8474 | 0.3329 | 0.9017 | 0.8038 | 0.8405 | 0.9357 | 0.8417 | 0.6701 |
| Std. Error |  |  |  |  |  |  |  |  |
| Y0 | 0.02527 | 0.03849 | 0.01835 | 0.02037 | 0.02294 | 0.03224 | 0.02625 | 0.02802 |
| Plateau | 0.006448 | 0.01448 | 0.004094 | 0.007067 | 0.005005 | 0.004843 | 0.00564 | 0.007901 |
| K | 0.000843 | 0.003078 | 0.0005928 | 0.0006812 | 0.0007993 | 0.001125 | 0.0009171 | 0.00116 |
| Span | 0.02315 | 0.0336 | 0.01704 | 0.01798 | 0.02135 | 0.03085 | 0.02447 | 0.02539 |
| 95% Confidence Intervals |  |  |  |  |  |  |  |  |
| Y0 | -0.04013 to | -0.06389 to | -0.08638 to - | -0.01760 to | -0.05608 to | -0.1463 to - | -0.05606 to | -0.05753 to |
|  | 0.05903 | 0.08716 | 0.01436 | 0.06236 | 0.03393 | 0.01982 | 0.04698 | 0.05244 |
| Plateau | 0.8442 to<br>0.8695 | 0.3161 to<br>0.3730 | 0.8433 to<br>0.8594 | 0.8123 to<br>0.8401 | 0.8196 to<br>0.8392 | 0.8431 to<br>0.8621 | 0.8260 to<br>0.8482 | 0.6521 to<br>0.6831 |
| K | 0.01638 to<br>0.01969 | 0.008330 to<br>0.02041 | 0.01844 to<br>0.02077 | 0.01371 to<br>0.01638 | 0.01832 to<br>0.02145 | 0.02324 to<br>0.02765 | 0.01828 to<br>0.02188 | 0.01470 to<br>0.01925 |
| Tau | 50.80 to<br>61.05 | 49.00 to<br>120.0 | 48.15 to<br>54.22 | 61.05 to<br>72.96 | 46.61 to<br>54.59 | 36.17 to<br>43.04 | 45.70 to<br>54.69 | 51.95 to<br>68.03 |
| Half-time | 35.21 to<br>42.32 | 33.96 to<br>83.21 | 33.38 to<br>37.59 | 42.32 to<br>50.57 | 32.31 to<br>37.84 | 25.07 to<br>29.83 | 31.67 to<br>37.91 | 36.01 to<br>47.15 |
| Span | 0.8020 to<br>0.8928 | 0.2670 to<br>0.3988 | 0.8683 to<br>0.9351 | 0.7685 to<br>0.8391 | 0.7986 to<br>0.8824 | 0.8751 to<br>0.9962 | 0.7936 to<br>0.8897 | 0.6203 to<br>0.7199 |
| Goodness of Fit |  |  |  |  |  |  |  |  |
| Degrees of Freedom | 1149 | 1149 | 1053 | 1053 | 1149 | 1149 | 957 | 1149 |
| R square | 0.6867 | 0.1122 | 0.8441 | 0.7471 | 0.7357 | 0.6725 | 0.7202 | 0.5177 |
| Absolute Sum of Squares | 15.39 | 47.97 | 6.027 | 10.69 | 10.97 | 14.27 | 9.822 | 20.59 |
| Sy.x | 0.1157 | 0.2043 | 0.07566 | 0.1007 | 0.09771 | 0.1114 | 0.1013 | 0.1339 |

**Table 3.** FRAP analysis deletion mutants.

| | WT | ΔPE | ΔABD | PE-only | ΔPS<br>(not in figures) |
| --- | --- | --- | --- | --- | --- |
| One-phase association |  |  |  |  |  |
| Best-fit values |  |  |  |  |  |
| Y0 | -0.05615 | -0.01408 | -0.1411 | -2.811 | -0.03120 |
| Plateau | 0.7485 | 0.7320 | 0.8718 | 0.8699 | 0.7179 |
| K | 0.01662 | 0.01220 | 0.02823 | 0.0004781 | 0.01296 |
| Tau | 60.18 | 81.94 | 35.43 | 2091 | 77.13 |
| Half-time (seconds) | 41.71 | 56.80 | 24.56 | 1450(msec) | 53.47 |
| Span | 0.8046 | 0.7461 | 1.013 | 3.681 | 0.7491 |
| Std. Error |  |  |  |  |  |
| Y0 | 0.01209 | 0.01963 | 0.02240 | 0.3488 | 0.02405 |
| Plateau | 0.003295 | 0.009428 | 0.002599 | 0.002656 | 0.01029 |
| K | 0.0003987 | 0.0006632 | 0.0007211 | 2.293e-005 | 0.0008132 |
| Span | 0.01091 | 0.01635 | 0.02164 | 0.3483 | 0.02037 |
| 95% Confidence Intervals |  |  |  |  |  |
| Y0 | -0.07986 to -0.03244 | -0.05258 to 0.02442 | -0.1850 to -0.09716 | -3.495 to -2.126 | -0.07838 to 0.01598 |
| Plateau | 0.7420 to 0.7549 | 0.7135 to 0.7505 | 0.8667 to 0.8769 | 0.8647 to 0.8751 | 0.6977 to 0.7380 |
| K | 0.01584 to 0.01740 | 0.01090 to 0.01350 | 0.02681 to 0.02964 | 0.0004332 to 0.0005231 | 0.01137 to 0.01456 |
| Tau | 57.48 to 63.15 | 74.05 to 91.72 | 33.74 to 37.30 | 1912 to 2309 | 68.68 to 87.96 |
| Half-time | 39.84 to 43.77 | 51.33 to 63.57 | 23.38 to 25.85 | 1325 to 1600(msec) | 47.61 to 60.97 |
| Span | 0.7832 to 0.8260 | 0.7140 to 0.7781 | 0.9704 to 1.055 | 2.997 to 4.364 | 0.7091 to 0.7890 |
| Goodness of Fit |  |  |  |  |  |
| Degrees of Freedom | 2205 | 1437 | 1341 | 1533 | 1149 |
| R square | 0.8218 | 0.6519 | 0.8362 | 0.5420 | 0.6214 |
| Absolute Sum of Squares | 12.62 | 20.83 | 6.211 | 14.52 | 18.69 |
| Sy.x | 0.07565 | 0.1204 | 0.06806 | 0.09732 | 0.1275 |

## Methods

### No statistical methods were used to predetermine sample size. Cell culture and media

Immortalized mouse embryonic fibroblasts (MEFs) derived from RPTP α+/+ murine background (Su, Muranjan and Sap, 1999) were grown in complete media composed by DMEM (Lonza) supplemented with 10% Fetal Bovine Serum South American (FBS SA, Euroclone) and 2mM Glutamine at 37°C and 5% CO_2_. Cells density never exceed 70% confluency to favor single-cell analysis instead of tissue-like behavior. For imaging experiments, MEFs were seeded on borosilicate glass coverslips of 1½ thickness (Corning) or Nunc Glass Base Dishes (Thermo Scientific) coated with sterile 10 µg/ml fibronectin (Sigma-Aldrich). During live imaging experiments, the culturing media was exchanged 30 minutes before experiments in serum-2+ deprived Ca -buffered RINGER solution (see Resource Table for buffer composition) to avoid Ca 2+ 2+ /Mg withdrawal shocks and avoid phenol-red background contamination during laser excitation. For experiments with AP2σ plasmid, RINGER solution was supplemented with 10% FCS. Other cell lines were obtained from IFOM Cell Bank and cultured as follow: NIH3T3, HS27, RPMI, A-431 were cultured in DMEM (Lonza) supplemented with 10% Fetal Bovine Serum South American (FBS SA, Invitrogen). For MCF-10A, DMEM-Ham’s F12 (Biowest VWR) was supplemented with 5% Horse Serum (Life Technologies), 10ug/ml Insuline, 20ng/ml EGF, 500ng/ml Hydrocortisone, 100ng/ml cholera toxin, 2mM L-Glutamine; MDA-MB 231 were cultured in Leibovitz’s L15 (Biowest VWR) supplemented with 10% FBS and 2mM L-Glutamine; HeLa were cultured in DMEM, 10% FBS, 1mM Na Pyruvate, 0.1mM NEAA, 2mM L-Glutamine. HUVEC primary cells were cultured in all-in-one ready-to-use Endothelial Cell Growth Medium (Cell application Inc. Merck). All reagents specific supplier and identifier are listed in the Resource Table.

### Plasmids and transient transfections

Plasmids for the mammalian transient expression used in this study are listed in the Resource Table (supplemental information), describing the original source and identifier. Specifically, human βII-Spectrin (gene ID: NM_003128) wild type construct was cloned into the pEGFP-C3 backbone (Clonetech) between HindIII and SacII restriction sites, with the fluorescent tag at the N-terminus interspaced by an additional Δ, residues 1776-1907 were deleted from full-length βII-Spectrin. The same peptide was cloned in frame with GFP into the pEGFP-N1 backbone to generate GFP-PE domain only. For the generation of GFP-βII-Spectrin-Δ PS, residues 421-530 were deleted from full-length βII-Spectrin. For the generation of GFP-βII-Spectrin-Δ D, residues 280-2364 were amplified from full-length GFP-βII-Spectrin, and cloned into pEGFP-C3 backbone between HindIII and SacII restriction sites. Other plasmids were purchased or obtained from external sources listed in the Resource Table. Transient expression was obtained by electroporation performed 24 hours before the experiment using the Neon electroporation system (Thermo Fisher 6 Scientific). For each transfection, 1×10 cells were trypsinized, washed once with PBS and mixed with a total of 10 μg recombinant DNA in electroporation buffer R (Invitrogen). Cells were singularly electroshocked at 1600mV for 20 msec by placing the electroporation tip into the column filled by E2 buffer (following manufacturer specifications). After the shock, cells were immediately seeded onto tissue-culture grade plastic dishes replenished with complete culturing media and let recover at least 12 hours.

### CONFOCAL and TIRF Microscopy

Confocal microscopy was performed on a Leica TCS SP8 laser scanning confocal microscope mounted on a Leica DMi 8 inverted microscope, equipped with a motorized stage and controlled by the software Leica Application Suite X (LASX) ver. 3.5.2.18963. For image acquisition, we used a HC PL APO CS2 63X/1,40 oil immersion objective. DIC, Epi-Fluorescence and Total Internal Reflection Fluorescence microscopy (TIRFM) of fixed specimens, live time-lapse of spreading cells, drug treatments, osmotic shocks, cell stretching (EPI mode only) and cell compression were performed on a Leica AM TIRF MC system. Two different objectives have been used: HCX PL APO 63X/1.47NA oil immersion and HCX PL APO 100X/1.47NA oil immersion. The lasers used for fluorochromes excitation were 488nm, 561nm, 635nm. A specific dichroic and emission filters for each wavelength have been used. The microscope was controlled by Leica Application Suite AF software (Ver. 2.6.1.7314) and images were acquired with an Andor iXon DU-8285_VP camera. For live imaging experiments, environmental conditions were maintained thanks to an Okolab temperature and CO_2_control system if needed.

### Micropatterning by Quartz Mask

Borosilicate glass coverslips (Corning) were washed with 70% Ethanol, airy-dried, activated with plasma cleaner (Harrick Plasma) for few minutes and incubated with 0.1 mg/ml PLL-PEG for 1 hour at room temperature. A quartz mask (Delta mask B.V.) wash washed with 70% Ethanol and activated under UV light using a UV lamp (UVO Cleaner, Jelight). PEGylated coverslips were put on the desired pattern in the mask, which then was illuminated under UV light for 7 minutes. Patterned coverslips were coated with 5-25 μg/ml fibronectin for 1 hour at room temperature, washed several times with PBS and 1-5×104 RPTP cells were seeded and cultured at 37 °C in the same media described before.seeded and cultured at 37 °C in the same media described before.

### Micropatterning by PDMS Stamping

Polydimethylsiloxane (Sylgard 184 silicone elastomere kit, Dow Corning) was prepared by mixing its two components in 1:10 ratio and degassed. It was then poured on the mold, degassed again and cured over night at 6 °C. PDMS was peeled of the molds and plasma cleaned to make the surface hydrophilic. The stamps were then coated with 10-30 μ fibronectin for 30 minutes at room temperature. Excess of fibronectin was airy-dried and the stamps were gently pressed onto the previously silanized borosilicate glass bottom dishes or coverslips for 1 minute and then carefully removed. Silanization was achieved by pouring a methanol solution containing 0.16% v/v silane (Sigma-Aldrich) for 1 hour, followed by three washes in pure methanol. To passivate the surface of the non-fibronectin coated glass, the surface was treated with a solution of 0.5% PEG-PLL (Ruixibio) for 1 hour in order to avoid cell attachment on the4 unstamped area. After the incubation, the dishes were rinsed several times with PBS and 1-5×10 were seeded and cultured at 37 °C in the same media described before.

### Immunofluorescences

The antibodies used in this study were the following: mouse anti-βII-spectrin (dilution 1:200, BD-bioscience), rabbit anti βII-spectrin (1:200, Abcam), mouse anti-αII-spectrin (1:200, Invitrogen), rabbit anti-β-actin (1:100, Cell Signaling) and mouse anti-Clathrin Heavy Chain (1:500, Thermo Fisher, clone X22). Before fixation, cells were seeded on 10 μ ml fibronectin-coated coverslips/glass-bottom dishes. Cell lines g/ were fixed in 4% paraformaldehyde in PBS for 10 minutes and then neutralized using 10 mM NH_4_Cl in PBS for 10 minutes. Alternatively, fixation was performed in ice-cold methanol for 2 minutes at −20°C when cells were immunostained by anti-β-actin antibody. Cells were subsequently washed three times with PBS (5 minutes each), permeabilized for 2-5 minutes using PBS containing 0.1% Triton X-100 and blocked with 3-5% BSA for 10 minutes at room temperature. Then, cells were incubated with primary antibody for 1 hour at room temperature or over-night at 4°C. After 3 washing steps in PBS, cells were incubated with Alexa 488/647-conjugated goat anti-mouse or anti-rabbit (1:100-1:400, Thermo Fischer) and Alexa 488/546-conjugated phalloidin (1:200, Sigma-Aldrich) for 1 hour at room temperature. After three washes in PBS of 5 minutes each, cells were mounted with glycerol (for confocal microscopy) or in PBS (for TIRF microscopy) and stored at °C. All primary antibodies and fluorophore-conjugated secondary antibodies, source and identifier are listed in Resource Table. For intensity correlation analysis (Pearson’s coefficient and scatter plot), the plug-in JACoP was used (Bolte and Cordelieres, 2006) and plotted using R ggplot2 package.

### Western Blotting

For western blot, cells seeded on plastic tissue culture grade dishes were lysate directly on the plate by adding modified Sample buffer composed by Tris-HCl 135mM (pH 6.8), Sodium dodecyl sulphate (SDS) 5%, Urea 3.5M, NP-40 2.3%, β-mercaptoethanol 4.5%, glycerol 4% and traces of bromophenol blue. Cells were normalized by seeding density since this lysis buffer does not allow total protein content measurements, but prevent membrane-bound proteins from being degraded during trypsinization. An equal volume of samples was then loaded on 12-8% SDS-PAGE gels and transferred to nitrocellulose membrane (Amersham GE-Healthcare). Membranes were blocked in 5% milk-TBS with 0.1-0.3% Tween-20 for 1 hour at room temperature, then incubated overnight at 4° with primary antibodies (mouse anti-βII-spectrin 1:2000 dilution (BD-bioscience), rabbit anti-βII-spectrin 1:2000 dilution (Abcam), mouse anti-β-tubulin dilution 1:5000 (Sigma-Aldrich) and 1 hour at room temperature with HRP-conjugated secondary antibodies (BioRad). Proteins were detected with ECL Western blotting reagents (Amersham GE-Healthcare), by the digital Chemidoc XRS+ run by the software Image Lab (Biorad).

### Fixed Spreading assay

MEFs were seeded on 10 μg/ml fibronectin-coated coverslips/glass-bottom dishes in complete media, as5 previously described. A total of 10 cells were seeded and fixed at different time points (between 5 and 20 minutes after seeding) by 4% paraformaldehyde diluted in PBS for 10 minutes. Before detergent permeabilization for immunostaining, fixed cells were incubated with membrane dye FM4-64 FX (Thermo Scientific) according to manufacturer specification (2.5-5 μg/ml). A second fixation step was applied to preserve cell membrane staining by the fixable dye and avoid diffusion after a subsequent permeabilization step in 0.1% Triton-X100 (1-2 minutes). Immunostaining for βII-spectrin and F-actin (phalloidin) was performed as previously described for TIRFM investigation. From the raw images, the signal background was subtracted, while edge-preserving filters were applied to the FM4-64 fluorescence signal to generate a binary mask of the projected cell area in the TIRF plane. The Fiji built-in “Analyze particle” tool was then used to extract the projected area value as well as the integrated density of the two immunostainings (βII-spectrin and F-Actin). Linear regression analysis between the three parameters was then performed by R software, and raw data points plotted by the software Graphpad Prism.

### Live Spreading assay

The spreading assay was performed on custom-designed 2-way aluminium slides, sealed on the two planar faces by 22×22 mm glass coverslips welded by high vacuum grease (Sigma-Aldrich). Coverslips were previously acid-washed with a 20% HNO_3_solution for 2 hours at room temperature, followed by a final wash in pure acetone before being dried and coated with 10 µg/ml fibronectin for 1 hour at 37°C. The chamber was rinsed with 1x RINGER solution and incubated on the microscope stage at 37°. The dual glass surfaces allow simultaneous fluorescence and DIC illumination during media addition/exchange. MEFs were transfected 24 hours before imaging with the opportune plasmid combinations. Before the experiment, cells were trypsinized, centrifuged 5 minutes at 1200 rpm, washed once with PBS and serum-starved in suspension for 30 minutes at 37° in CO_2_-independent 1X RINGER solution. Suspended cells were thereafter4 kept at room temperature up to 3 hours. For each time-lapse, 1-5×10 cells were fluxed into the imaging chamber to optimally observe single-cell spreading; for this reason cell aggregates or debris were carefully avoided during imaging. The time-lapse started after a positively double-transfected cell engaged with the fibronectin-coated surface; cells were then followed for 15-20 minutes at time rates of 2-5 seconds/frame. For experiments in presence of myosin II inhibitor blebbistatin (Sigma-Aldrich), cells were suspended in 1x RINGER solution supplemented with blebbistatin at 50μM final concentration; imaging chambers were filled with the same 1x RINGER solution supplemented with blebbistatin to avoid rebound effects upon injection of cell suspension. The correct cell behaviour was monitored by DIC acquisition, in particular by focusing on cell integrity, isotropic spreading in P1, lamellipodia formation and buckling. The fluorescent channels were analysed as described in the next sections for cell edge and body behaviour.

### Spectrin and actin intensities at the cell edge

For cell edge analysis a custom macro for Fiji was written. The signal background was subtracted, while edge-preserving filters were applied to the actin fluorescence signal to generate a binary mask of the growing cell area over time. The centroid of the cell was used as a reference point to identify each angle between 0⁰ and 359⁰ on the cell outer circumference. From the edge, the signal was eroded by 25 pixels (≈3.2µm at the resolution of ≈130nm/pixel) and mean intensity of the fluorescent signals were computed into the final kymographs composed of 360 pixels on the x-axis, one for each angle, while the y-axis represents the total number of frames. Speed of the cell edge was extrapolated from the distance variation in pixel between the outer edge and the cell centroid, at known pixel size and time frame. Values were considered positive when the edge moved away from the centroid and negative when it moved closer. For comparisons between independent cells, actin and spectrin intensity measurements were normalized to 1 at null speed. Results were analyzed and plotted with R and the ggplot2 R package.

### Correlated Spectrin and actin flow velocity analysis (PIV)

Correlation between speed and directionality of the two fluorescent channels have been performed on the same live TIRFM datasets during the spreading analyzed for cell edge dynamics. The signal background was subtracted. Particle Image Velocimetry (PIV) was performed independently on single fluorescent channels by a custom macro in Fiji, excluding the portion of the cell close to the edge (50 pixels) and the frames corresponding to the initial spreading phase P1 from the analysis (50 frames). The resulting vector fields of the two channels (i.e. RFP-actin and GFP-βII-spectrin) mapped both the speed magnitude and directionality (angle) of the channels independently. To identify synchronous angular and speed correlations between the two channels, their correlation was binarized by giving a value of 1 to areas where the speeds of the two channels were within +/-50% of each other, and where the directions shared the same quadrant (i.e. when the difference of angle was less than π/2). Areas that did not meet these criteria were assigned a value of 0. The fraction of correlated flow velocities was then given as the ratio between the area covered by correlated velocities and the area of the cell after excluding the edge portion. The area of correlated motion was calculated for each binarized frame by the “Analyze particle” tool, and by randomly applying the Watershed algorithm to segment neighbour areas with no morphological continuity. The same parameters were blindly applied to the untreated and blebbistatin-treated time-lapses to avoid bias. Results were analyzed and plotted with R and the ggplot2 R package.

### Cytoskeletal Drug Perturbation and osmotic shocks

MEFs were transfected 24 hours before imaging with the opportune pair of constructs as previously described. Before imaging, cells were trypsinized, seeded on glass coverslip coated with 10 µg/ml fibronectin (1×10^3^ cells) and allowed to attach to the substrate for 1 hour in complete media. The media was replaced with CO_2_-independent 1x RINGER solution and mounted on a 2-way imaging chamber that allows on-stage media exchange as previously described for the spreading assay. Time-lapse consisted of an initial phase of 5 minutes where sufficient frames were acquired at steady-state (internal control); the media was then replaced by 1x RINGER solution supplemented with the opportune cytoskeletal impairing drugs. Addition of media exceeded the volume of the imaging chamber to avoid dilution of final drug concentrations and rebound effect. Specifically, 1µM Latrunculin A, 10µM Blebbistatin, 5 µM Nocodazole and 5mM Diamide were singularly used (Sigma-Aldrich). Perturbed cells were then imaged for 30 minutes at 1-5 minute/frame. Intensity calculations were carried out in Fiji by subtracting the background and creating a dynamic binary mask of the GFP-βII-Spectrin signal; the built-in “Analyze particle” tool was applied to obtain mean fluorescence intensity values at different time points. Only untreated, 5 and 30 minutes after treatment time points were plotted using the software Graphpad Prism.

For experiments that required osmotic shocks, MEFs cells were treated following the same procedure described earlier for cytoskeletal perturbations. Time-lapse was obtained at higher temporal resolution (2-5 second/frame). At given time points (every 3-5 minutes depending on the experiment), the media was replaced with hypotonic 0.5x or 0.1x RINGER solution, exceeding the volume of the imaging chamber to avoid incomplete media exchange. Mean intensity calculation was done in Fiji by subtracting the background and creating a dynamic binary mask to derive changes in fluorescence intensity over time. In the case of ratio measurements between the two fluorescent channels, mean intensities of the first two frames of the meaningful channels were averaged and arithmetically matched to obtain the initial ratio value of 1 (non-stoichiometric). Indeed, the purpose of these experiments was to calculate the fluctuation in content more than a stoichiometric measurement between the two fluorescent proteins. Intensity data were then averaged, analyzed and plotted using the software Graphpad Prism.

### Cell stretching

Cell stretching experiments were performed using an automated cell stretching dish (International patent: WO 2018/149795 A1). The components of the cell stretching dish were designed using SolidWorks software and 3D printed using a stereolithography-based 3D printer (Form 2, Formlabs) coupled with an autoclavable and biocompatible dental resin (Dental SG resin, Formlabs). The printed parts were rinsed in isopropyl alcohol for 5 minutes to remove any uncured resin from their surface, and then post-cured in a UV box to finalize the polymerization process and stabilize mechanical properties. The printed parts were then polished and assembled to create the lower (cell chamber) and the upper portion (aperture driver) characterizing the stretching dish. Before the experiments, the components of the lower portion were autoclaved to be sterilized and assembled to clamp a deformable silicone membrane (thickness 0.0051, SMI), thus creating a cell culture chamber. The dish was coated with fibronectin (10 µg/ml) and incubated at 37 °C for 1 hour. A total of 10^5^ cells were seeded in the cell chamber and let spread for 1-2 hours at 37 °C. Before starting the imaging, the whole-cell stretching dish was assembled by connecting the upper portion, consisting in the driving unit, to the cell culture chamber. Biaxial stretching was applied to all the samples under investigation. The EPI-fluorescence mode of the Leica AM TIRF MC system was used. Due to technical limitations such as re-focusing and re-centering of the cell under investigation, after each 5% step increase in the biaxial stretch obtained by the software-controlled motorized device, a single frame in the two fluorescent channels was recorded.

### Cell compression

Cell compression experiments were performed using a cell compression device (International patent: WO 2019/086702 A1) capable of applying dynamic compression stress to single cells. The main components of the compression device were designed using SolidWorks software and 3D printed using a stereolithography-based 3D printer (Form 2, Formlabs), following the same procedure described in for the cell stretching device. The cell compression device consists of an air chamber connected to an air pressure regulator. Before the experiments, a Polydimethylsiloxane (PDMS, Sylgard 184) piston was microfabricated to have circular pillars (200 µm in diameter) on its surface, and attached to a deformable silicone membrane (thickness 0.0051, SMI) through plasma bonding. The membrane with the piston was then clamped to the air chamber. The assembled cell compression device was connected to the air pressure regulator and then locked to the cell culture dish (seeded with cells) through mechanical ribs. In particular, 5 a total of 10 cells were seeded on 27mm Ø Nunc Glass Base Dishes (Thermo Scientific), exchanged to 1x RINGER solution 30 minutes before imaging and the original Petri lid replaced by the compression device. A dynamic compressive load was applied to cells by increasing the air pressure inside the compression device through the pressure regulator, thus controlling the movement of the membrane and the piston to compress the cells underneath. DIC and TIRF illumination were used to monitor cell reaction at 5 seconds/frame rate. ROIs to measure fluorescence intensity fluctuation was drawn in Fiji, while the DIC was used to monitor the engagement of the piston with the cell roof. As the compression strain increased, the cell became flatter and DIC imaging decreased its contrast in physiologically flat fibroblasts. Due to cell height variation, the device does not allow precise absolute read-out of the pressure applied to cells, being cell deformation a non-controllable variable. For this reason, two different approaches were applied. The first one was intended to cause maximal cell response: compression pressure was slowly increased until bleb formation was observed. For i-bleb/cortex ratio, mean fluorescence intensities in the two channels were obtained in the projected bleb area, and divide by the mean fluorescence intensities in the adjacent cortical region of similar area. The second approach was designed to better control the applied strain: the initial compression was thus set at the first value required to engage the piston with the top of the cell as monitored by DIC. This first step hardly caused a reaction monitorable by TIRF at basal cell level but allowed consistency between independent experiments, for this reason six different cycles of compression (2 minutes) and relaxation (2 minutes) were performed at increasing pressure, as schematized in D; the 2 minutes compression step duration was chosen to allow adaptive mechanisms to occur and avoid long-term detrimental effects on cell integrity. Intensity calculations were carried out in Fiji by subtracting the background and drawing ROIs across the perinuclear rim formed during compression. Pre-stretch mean intensity for each single compressive events was divided by maximal mean intensity registered during the subsequent 2 minutes compressive step. Data were plotted using the software Graphpad Prism.

### Frap experiments

MEFs cells expressing GFP-βII-spectrin constructs were imaged 24 hours after transfection with a Confocal Spinning disk microscope (Olympus) equipped with iXon 897 Ultra camera (ANDOR) and a FRAP module furnished with a 405nm laser. The environmental control was maintained with an OKOlab incubator. Images were acquired using a 100x/1.35Sil silicone oil immersion objective. MEFs were trypsinized and seeded on glass-bottom dishes (Matek, Sigma-Aldrich) coated with 10 µg/ml fibronectin. Before imaging, CO_2_-independent media without phenol-red was exchanged. Squared Regions Of Interest of 5×5µm length were photo-bleached with the 405 nm laser at 50% intensity and post-bleach images were followed with 15 to 20% laser intensity for 100 frames (1 frame every 5 seconds for full-length and truncated GFP-βII-spectrin constructs and 0.5 seconds for PE-only). FRAP data were analyzed and curves fitted to the one-exponential recovery equations (one-phase association) by the software Graphpad Prism:

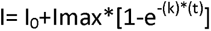

Where I represents the relative intensity compared to the pre-bleach value, k the association rate, and t the half time recovery expressed in seconds.

For the dual FRAP assay, cells were seeded on a fibronectin-coated glass coverslip and mounted on the 2-way imaging chamber that allows on-stage media exchange as previously described for spreading assay and osmotic shocks. The first FRAP measurements were conducted in serum-free 1x Ringer solution; at completion, the media was exchanged and independently supplemented with the cytoskeletal drugs at the concentration used before. Specifically, 1 µM Latrunculin A, 10 µM Blebbistatin, 5 µM Nocodazole and 5 mM Diamide (Sigma-Aldrich). Cells were allowed to equilibrate with the new media for 5 minutes; afterwards, a second FRAP analysis was started on the same cell previously analyzed.

### Clathrin pits density maps

TIRFM images were acquired as previously described. For analysis of fixed specimens, 2×2 µm ROIs were selected by segmenting only discrete pits of random sizes, not overlapping with neighbouring structures (plaques were not considered for the analysis). All images were stacked in FIJI for pit size calculation. CHC/AP2 Images were up-scaled by a factor of 10, Yen auto-threshold applied to create a binary mask and particle size calculated by the “Analyze particle” tool in FIJI. ROIs were divided at this point according to the size that took into account the diffraction limits of TIRFM (<300nm^2^ and 300-500nm^2^); particle bigger that 2 500nm were not considered informative. Raw images of the two clusters were then independently stacked and z-projected for median intensity values. Gaussian blur filter (1 pixel radius) and a scale factor of 10 were applied to the projected images to homogenize the signals. Scaled-projected images of the two particle-size groups were then matched for signal intensities between corresponding channels. Normalized radial plots were generated by the Radial Profile plugin (FIJI) on the final 2×2 µm projected images.

For live imaging datasets, Unsharp Mask (1 pixel radius, 0.6 weight) and Gaussian blur filter (1 pixel radius) were applied to the raw images before ROIs selection. No additional filters were applied to the final projected images. Similar filtering strategy was applied in live datasets during osmotic shocks.

### RNA EXTRACTION AND qPCR ANALYSIS

Cells were cultured as previously described. At least 2×10^2^ cells were lysed and RNA was extracted with the RNAeasy Mini Kit (Qiagen) following manufacturer specifications. 1 µg of RNA was retrotranscribed using “qScript cDNA Synthesis kit” (Quantabio). For gene expression analysis, 5ng of cDNA was amplified (in triplicate) in a reaction volume of 10 μL containing the following reagents: 5 μl of TaqMan® Fast Advanced Master Mix, 0.5 μl of TaqMan Gene expression assay 20x (Thermo Fisher). The entire process (retrotranscription, gene expression and data analysis) was performed by the qPCR-Service at Cogentech-Milano, following the ABI assay ID Database (Thermo Fischer). Therefore, only gene ID of the spectrin murine genes analyzed could be provided here (Mm01315345_m1, Mm01180701_m1, Mm01326617_m1, Mm00661691_m1, Mm01284057_m1, Mm01239117_m1). Murine SPTBN5 primers were custom designed (Fw: GGACGCCAGTGTTCACCAA Rev: GCCCCCTTGTAGCAGCTT) since were not implemented in the database. Real-time PCR was carried out on the 7500 Real-Time PCR System (Thermo Fisher), using a pre-PCR step of 20 seconds at 95°C, followed by 40 cycles of 1 second at 95°C and 20 seconds at 60°C. Samples were amplified with primers and probes for each target and for all the targets one NTC sample was run. Raw data (Ct) were analyzed with “Biogazelle qbase plus” software and the fold change was expressed as CNRQ (Calibrated Normalized Relative Quantity) with Standard Error (SE). Gapdh was used as references to normalize the data. Three independent experiments were then averaged and plotted using the software Graphdap Prism.

### Statistical analysis

All the graphs and plots are presented as mean ± SD, except the FRAP recovery curves that are presented as mean ± SEM. These experiments were further analysed by fitting mean values to an exponential equation, therefore the authors considered trivial to show the accuracy of the mean (better described by the definition of SEM) instead of the intrinsic variability between independent experiments. Half-time recovery data was then plotted as mean ± 95% confidence interval of the fitting. Statistical analysis by unpaired T-test was performed when the comparison between two experimental groups was required (i.e. normalized i-bleb/cortex ratio). One-way Anova analysis was performed when multiple experimental groups were present. One-way Anova analysis with multiple comparisons between groups was also performed in parallel, as shown for the quantification of maximal intensity at the cell body during the sequential compression protocol. All the experiments were performed in triplicate or more, as indicated in the figure legends.

## CONTACT FOR RESOURCE SHARING

Further information and requests for resources and reagents will be provided upon reasonable request. Inquiries should be addressed and fulfilled by the Lead Contact, Nils Gauthier.

## Resource Table

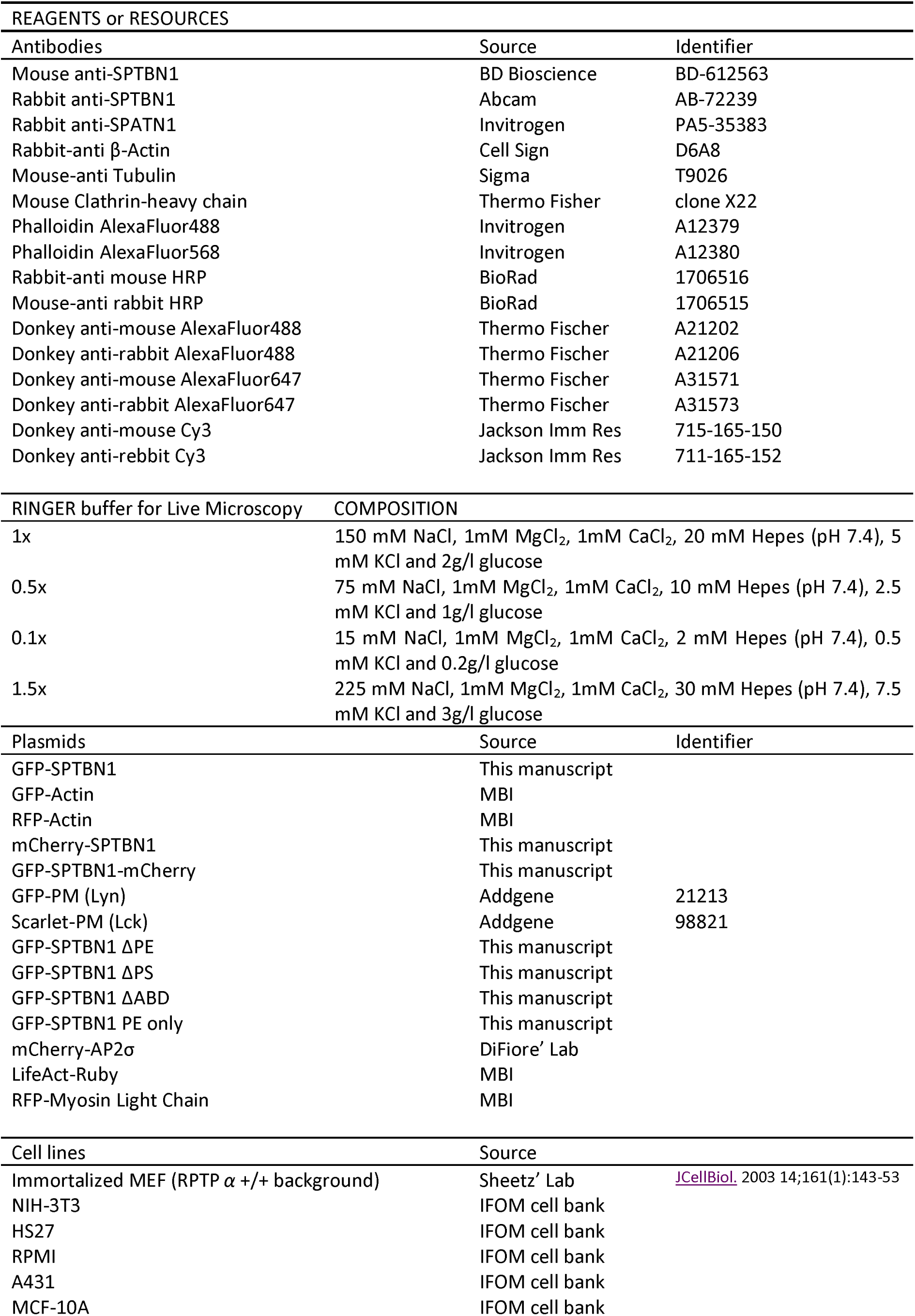
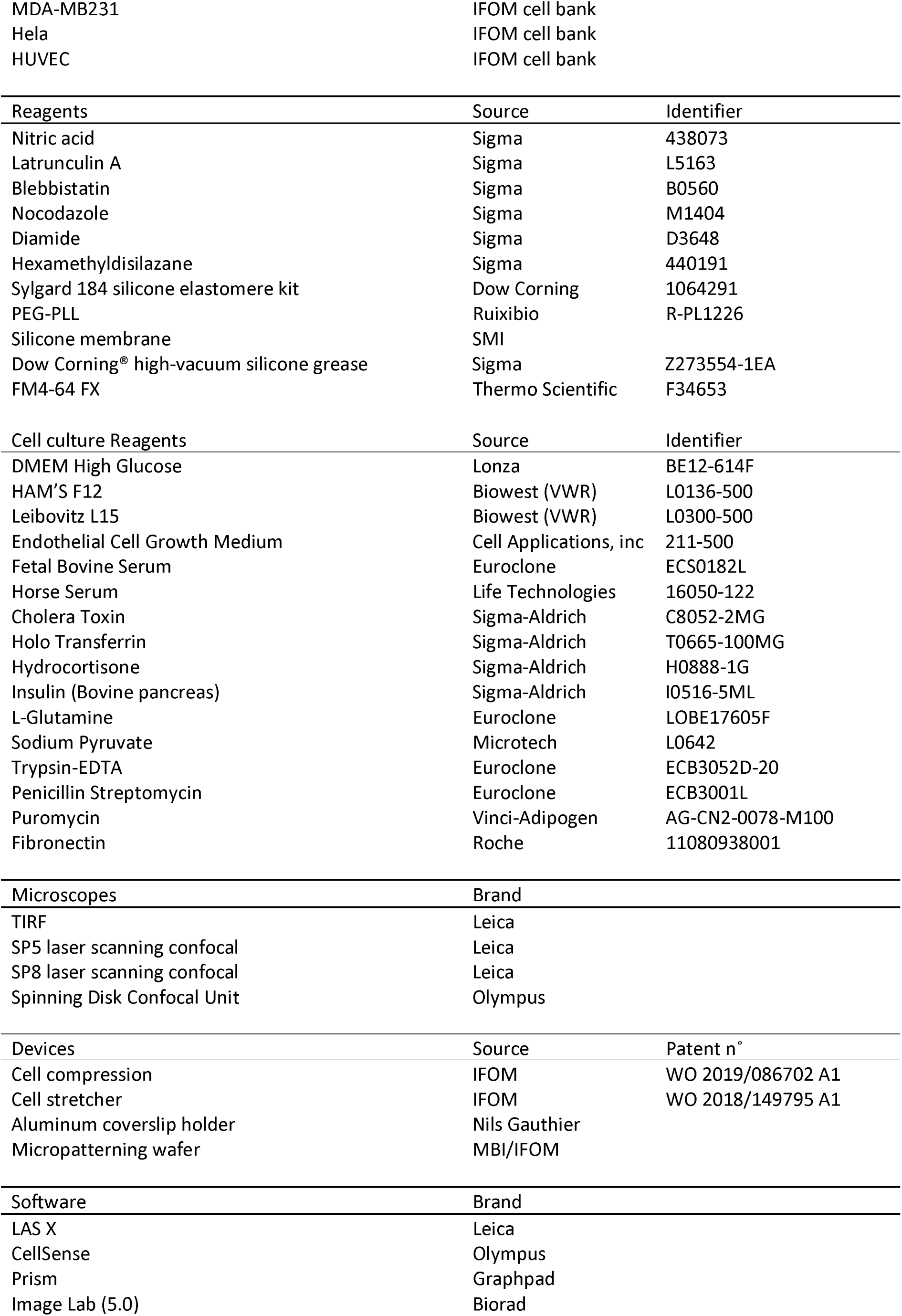
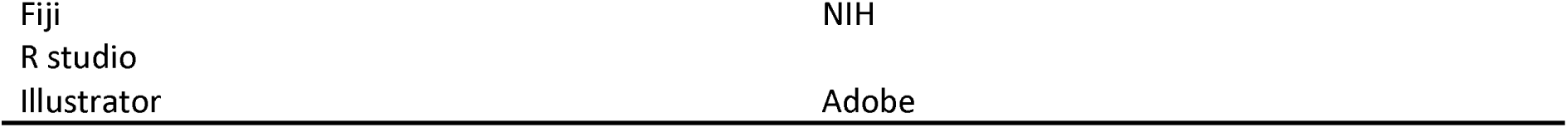

